# FXR inhibition reduces ACE2 expression, SARS-CoV-2 infection and may improve COVID-19 outcome

**DOI:** 10.1101/2021.06.06.446781

**Authors:** Teresa Brevini, Mailis Maes, Gwilym J. Webb, William T. H. Gelson, Sally Forrest, Petra Mlcochova, Scott Dillon, Sagar Varankar, Mahnaz Darvish-Damavandi, Victoria L. Mulcahy, Rhoda E. Kuc, Thomas L. Williams, Vasileios Galanakis, Marta Vila-Gonzalez, Olivia C. Tysoe, Daniele Muraro, Thomas W. M. Crozier, Johannes Bargehr, Sanjay Sinha, Sara S. Upponi, Lisa Swift, Kourosh Saeb-Parsy, Susan E. Davies, Thomas Marjot, Eleanor Barnes, Ansgar W. Lohse, Andrew M. Moon, A. Sidney Barritt, Ravindra K. Gupta, Stephen Baker, Anthony P. Davenport, Gareth Corbett, Simon J. A. Buczacki, Joo-Hyeon Lee, Paul Gibbs, Andrew J. Butler, Christopher J. E. Watson, George F. Mells, Gordon Dougan, Ludovic Vallier, Fotios Sampaziotis

**Affiliations:** Wellcome – MRC Cambridge Stem Cell Institute, Cambridge, UK; Cambridge Institute of Therapeutic Immunology & Infectious Disease (CITIID), Department of Medicine, University of Cambridge, Cambridge, UK; Cambridge Liver Unit, Cambridge University Hospitals NHS Foundation Trust, Cambridge, UK; Department of Medicine, University of Cambridge, Cambridge, UK; Nuffield Department of Surgical Sciences, Old Road Campus Research Building, Oxford, UK; Academic Department of Medical Genetics, University of Cambridge, Cambridge, UK; Experimental Medicine and Immunotherapeutics, University of Cambridge, Addenbrooke’s Hospital, Cambridge, UK; Department of Surgery, University of Cambridge and NIHR Cambridge Biomedical Research centre, Cambridge, UK; Wellcome Sanger Institute, Wellcome Genome Campus, Hinxton, Cambridge, UK; Division of Cardiovascular Medicine, University of Cambridge, Cambridge, UK; Department of Radiology, Cambridge University Hospitals NHS Foundation Trust, Cambridge, UK; Department of Surgery, Cambridge University Hospitals NHS Foundation Trust, Cambridge, UK; Department of Histopathology, Cambridge University Hospitals NHS Foundation Trust, Cambridge, UK; Oxford Liver Unit, Translational Gastroenterology Unit, Oxford University Hospitals NHS Foundation Trust, University of Oxford, Oxford, UK; Department of Medicine, University Medical Centre Hamburg-Eppendorf, Hamburg, Germany; Division of Gastroenterology and Hepatology, University of North Carolina, Chapel Hill, NC, USA; Cambridge University Hospitals NHS Foundation Trust, Cambridge, UK; Department of Physiology, Development and Neuroscience, University of Cambridge, Cambridge, UK; National Institute of Health Research (NIHR) Cambridge Biomedical Research Centre, and the NIHR Blood and Transplant Research Unit (BTRU) at the University of Cambridge in collaboration with Newcastle University and in partnership with NHS Blood and Transplant (NHSBT), Cambridge, UK

## Abstract

Prevention of SARS-CoV-2 entry in cells through the modulation of viral host receptors, such as ACE2, could represent a new therapeutic approach complementing vaccination. However, the mechanisms controlling ACE2 expression remain elusive. Here, we identify the farnesoid X receptor (FXR) as a direct regulator of ACE2 transcription in multiple COVID19-affected tissues, including the gastrointestinal and respiratory systems. We demonstrate that FXR antagonists, including the over-the-counter compound z-guggulsterone (ZGG) and the off-patent drug ursodeoxycholic acid (UDCA), downregulate ACE2 levels, and reduce susceptibility to SARS-CoV-2 infection in lung, cholangiocyte and gut organoids. We then show that therapeutic levels of UDCA downregulate ACE2 in human organs perfused *ex situ* and reduce SARS-CoV-2 infection *ex vivo*. Finally, we perform a retrospective study using registry data and identify a correlation between UDCA treatment and positive clinical outcomes following SARS-CoV-2 infection, including hospitalisation, ICU admission and death. In conclusion, we identify a novel function of FXR in controlling ACE2 expression and provide evidence that this approach could be beneficial for reducing SARS-CoV-2 infection, thereby paving the road for future clinical trials.

## Introduction

Vaccines for coronavirus disease 2019 (COVID-19) represent a major breakthrough against severe acute respiratory syndrome coronavirus 2 (SARS-CoV-2) infection. Nonetheless, they are not efficient against established disease and they are limited by variable efficacy^1^, the emergence of vaccine-resistant viral variants^2, 3^, cost^4^ and availability^5^. Treatment options are limited; while the drugs currently used, such as dexamethasone and remdesivir improve clinical outcome only in specific patient groups^6, 7^. Therefore, there is a pressing need for novel prophylactic and therapeutic agents. Viral host receptors represent ideal therapeutic targets^8, 9^, because they are essential for SARS-CoV-2 cellular entry and infection^10^. Among these, the angiotensin converting enzyme 2 (ACE2), is particularly appealing^11^. ACE2 is a transmembrane carboxypeptidase with a broad substrate specificity, including angiotensin II, which acts as the main receptor for SARS-CoV-2^12, 13^. It directly binds the spike protein of different coronaviruses, with a high affinity for SARS-CoV-2, rendering it indispensable for viral entry^14^. Accordingly, COVID-19 predominantly affects tissues expressing ACE2, such as the lungs, the cardiovascular system, the digestive tract and the biliary tree^15^. Modifying ACE2 expression could impede viral entry and protect against infection from SARS-CoV-2 and potentially other coronaviruses using the same receptor. Furthermore, because ACE2 is a host cell protein, its expression is not affected by mutations in the virus. Therefore, therapies modulating ACE2 expression may be effective against multiple SARS-CoV-2 variants and less prone to resistance^16^. However, the mechanisms controlling ACE2 expression remain elusive, while interspecies variation limits the use of animal models^17^. Here, we use human cholangiocyte organoids as a proof-of-principle system to demonstrate that the bile acid receptor farnesoid X receptor (FXR) controls ACE2 expression. We show that this mechanism applies in multiple SARS-CoV-2-affected tissues, including gastrointestinal and respiratory epithelia. We then demonstrate that FXR inhibition, with the approved drug ursodeoxycholic acid (UDCA) or the over-the-counter phytosteroid z-guggulsterone (ZGG), reduces ACE2 levels, conferring resistance to SARS-CoV-2 infection *in vitro*. We repeat our experiments in human organs perfused *ex-situ* and show that ‘systemic’ UDCA administration in circulating blood reduces ACE2 levels and viral infection. Subsequently, we measure ACE2 activity in the serum of patients before and after commencing UDCA for cholestatic disorders and observe a reduction in ACE2 activity; while we also validate that tissue from patients on UDCA exhibits lower ACE2 levels compared to tissue from organ donors not receiving UDCA. Finally, we interrogate an international registry cohort of patients with COVID-19 and chronic liver disease and describe a correlation between UDCA therapy and better clinical outcomes from COVID-19.

### Bile acids modulate ACE2 in cholangiocytes

To explore the mechanisms controlling ACE2 expression, we used cholangiocyte organoids (COs), as a proof-of-principle system. Our choice was based on a serendipitous finding from studying the involvement of the biliary tree in gastrointestinal (GI) COVID-19. To assess whether cholangiocytes are infected by SARS-CoV-2, we characterized the expression of viral entry receptors in the biliary epithelium using single-cell RNA sequencing (scRNAseq) (Extended Data Fig. 1a-c). We found that ACE2 was predominantly expressed in the gallbladder, compared to intrahepatic ducts (Wilcoxon Rank-Sum test P<4.5^-201^) (Extended Data Fig. 1b-d). We interpreted this result in the context of our recent work demonstrating that cholangiocytes are plastic, and in particular, that any cholangiocyte grown as an organoid *in vitro* can adopt a gallbladder identity after treatment with physiological levels of bile acids^18^. Therefore, we hypothesized that bile acids may also control the expression of ACE2. To explore this hypothesis, we interrogated the impact of chenodeoxycholic acid (CDCA), which is the main bile acid present in bile, on viral receptor levels in these organoids. We observed that CDCA upregulated ACE2 expression in cholangiocyte organoids to levels comparable to *in vivo* gallbladder cholangiocytes, regardless of the organoids’ region of origin (Extended Data Fig. 1a-c, 1e 2a-b). The relevance of this increased ACE2 expression for SARS-CoV-2 infection was illustrated first by identifying that the virus affects predominantly gallbladder cholangiocytes in the liver of COVID-19 patients (Extended Data Fig. 3a-b), which exhibit the highest ACE2 levels in the biliary tree; and then by demonstrating that cholangiocyte organoids treated with CDCA could be infected with SARS-CoV-2 *in vitro* (Extended Data Fig. 4a-c, 5a-m), produced infective viral progeny (Extended Data Fig. 4d), and appropriately upregulated the expression of innate immune genes and antiviral response markers (Extended Data Fig. 4e). Importantly, cholangiocyte organoids derived from any region of the biliary tree showed the same response to bile acid treatment resulting in overlapping transcriptional profiles (Extended Data Fig 1a-c, 1e); however, for consistency cholangiocyte organoids originating from the gallbladder (gallbladder cholangiocyte organoids - GCOs) were used for subsequent experiments. These results validated the role of cholangiocytes in COVID-19 and illustrated the value of cholangiocyte organoids as a platform to study SARS-CoV-2 infection; but most importantly, they identified that CDCA treatment increases ACE2 expression in these cells.

**Figure 1.**
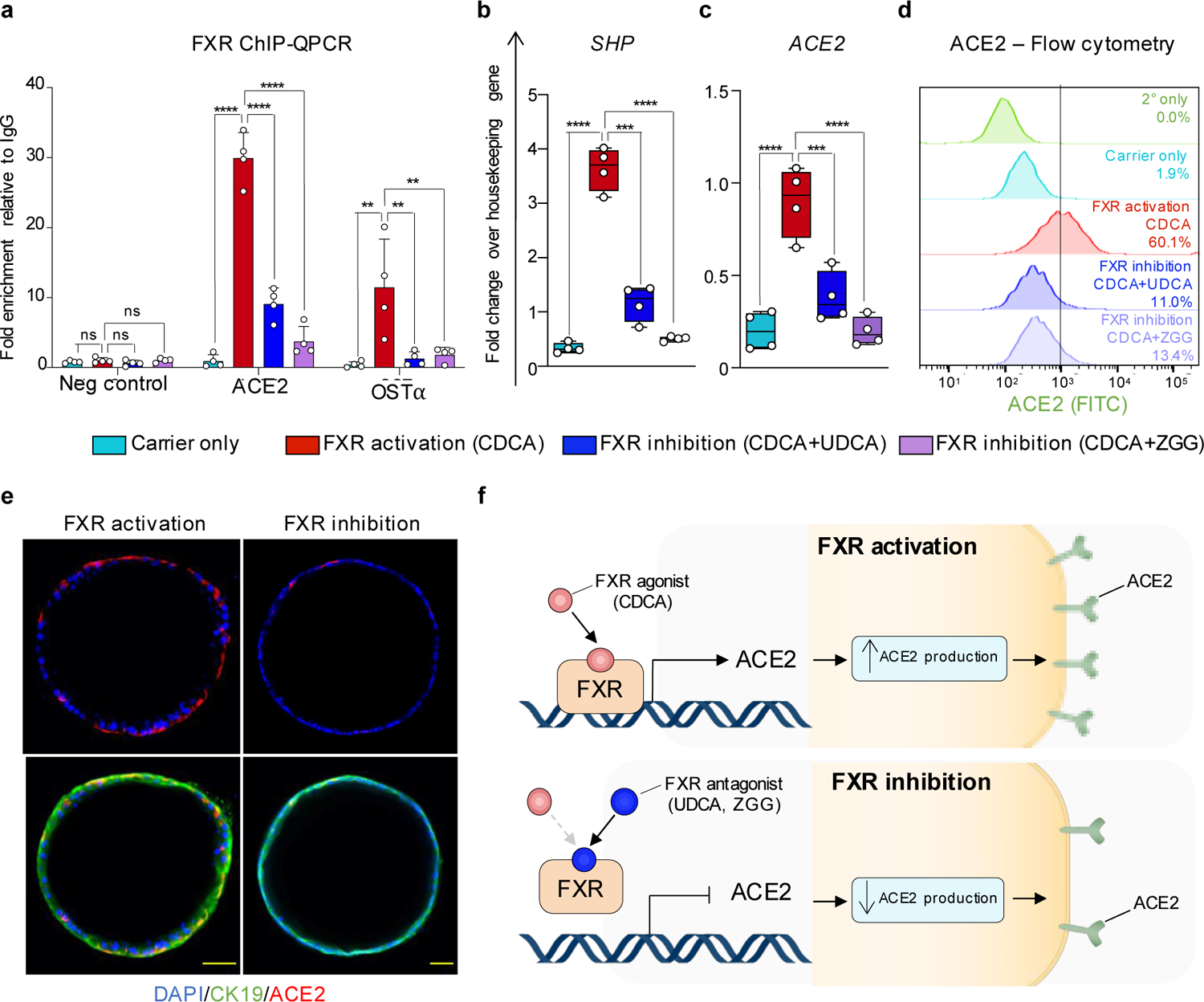
FXR controls ACE2 expression. (**a**) Chromatin immunoprecipitation followed by QPCR (ChIP-QPCR) on COs showing that the FXR agonist chenodeoxycholic acid (CDCA) promotes binding of FXR on the ACE2 promoter which is inhibited by FXR antagonists ursodeoxycholic acid (UDCA) and z-guggulsterone (ZGG). OSTα is used as positive control; n=4 independent experiments. (**b**) QPCR validating that CDCA, UDCA and ZGG respectively activate or inhibit FXR, as evidenced by corresponding changes in the expression of the FXR downstream effector SHP. (**c-e**) QPCR (**c**), flow cytometry (**d**) and immunofluorescence (**e**) showing changes in ACE2 levels upon modulation of FXR activity. Housekeeping gene, HMBS; n=4 independent experiments. One-way ANOVA with Dunnett’s correction for multiple comparisons; ns, non-significant; **, *P*<0.005; ***, *P*<0.001; ****, *P*<0.0001; center line, median; box, interquartile range (IQR); whiskers, range; error bars, standard deviation. Scale bars 50 μm. (**f**) Schematic representation of the suggested mechanism for FXR-mediated control of the viral receptor ACE2.

### FXR controls ACE2 in cholangiocytes

Since the bile acid CDCA is the most potent natural agonist of the bile acid receptor and transcription factor FXR^19, 20^, we hypothesized that CDCA could control ACE2 expression acting through FXR. To test this hypothesis, we first confirmed that FXR is expressed in gallbladder cholangiocytes *in vivo* and in GCOs *in vitro* (Extended Data Fig. 6a-c), and that it can be activated by CDCA treatment, as evidenced by the upregulation of its downstream target Small Heterodimer Partner (SHP) (Extended Data Fig. 6c). To assess whether FXR could bind the ACE2 gene and potentially control its transcriptional activity, we analysed the ACE2 promoter region and identified the presence of the FXR responsive element (FXR RE) IR-1. Accordingly, chromatin immunoprecipitation (ChIP-QPCR) confirmed that activated FXR binds on the ACE2 promoter (Fig. 1a). Conversely, FXR inhibition, using the FXR antagonist ZGG^21^ or the competitive antagonist UDCA^20, 22^, reduced FXR activity as evidenced by decreased SHP levels (Fig. 1b); decreased FXR presence on ACE2 promoter (Fig. 1a); and downregulated the expression of ACE2 at the transcript and protein levels (Fig. 1c-e). Considered together, these results demonstrate that FXR directly controls ACE2 expression in cholangiocytes (Fig. 1f).

### FXR controls ACE2 in multiple cell types

FXR is expressed in multiple cell types^20, 23–25^, and it can be activated by bile acids present in the systemic circulation^26^. Thus, ACE2 regulation through FXR may represent a general mechanism, extending beyond cholangiocytes. To explore this possibility, we repeated our experiments using primary organoids from key organs affected by COVID-19^27^, such as the lungs and the intestine. Importantly, the relevance of these platforms for studying SARS-CoV-2 infection has already been demonstrated^28, 29^. We first confirmed FXR expression in these tissues both *in vivo* and *in vitro* (Extended Data Fig. 6a-c). Subsequently, we showed that treatment with physiological concentrations of CDCA (10 μM)^26^ resulted in FXR activation, evidenced by upregulation of the FXR downstream target SHP (Extended Data Fig 6c) and increased ACE2 expression (Extended Data Fig. 7a-f). Conversely, FXR inhibition by UDCA or ZGG reduced ACE2 levels in primary airway and intestinal organoids (Extended Data Fig. 7a-f). These results confirm that FXR participates in the regulation of ACE2 expression in multiple cell types affected by COVID-19, suggesting that FXR-mediated control of ACE2 expression may represent a mechanism relevant for several organs (Fig. 1e).

### FXR inhibition limits SARS-CoV-2 infection *in vitro*

Our results show that FXR antagonists, including the clinically approved drug UDCA, used as first-line treatment in primary biliary cholangitis (PBC)^30^ and the over-the-counter drug ZGG, can reduce ACE2 expression in multiple cell types. To explore the relevance of this finding for COVID-19, we investigated whether ACE2 downregulation through FXR inhibition could reduce the susceptibility to SARS-CoV-2 infection *in vitro*. For this, we exposed gallbladder cholangiocyte, airway and intestinal organoids to physiological levels of CDCA, to simulate the baseline level of FXR activation present *in vivo*; and infected them with SARS-CoV-2 isolated from a patient’s naso-pharyngeal swab^45^ in absence or presence of the FXR antagonists UDCA or ZGG (Fig. 2a). FXR inhibition reduced viral infection in all three types of organoids (Fig 2b-e). These results confirm that FXR inhibition increases resistance to SARS-CoV-2 infection in multiple cell types *in vitro*.

**Figure 2.**
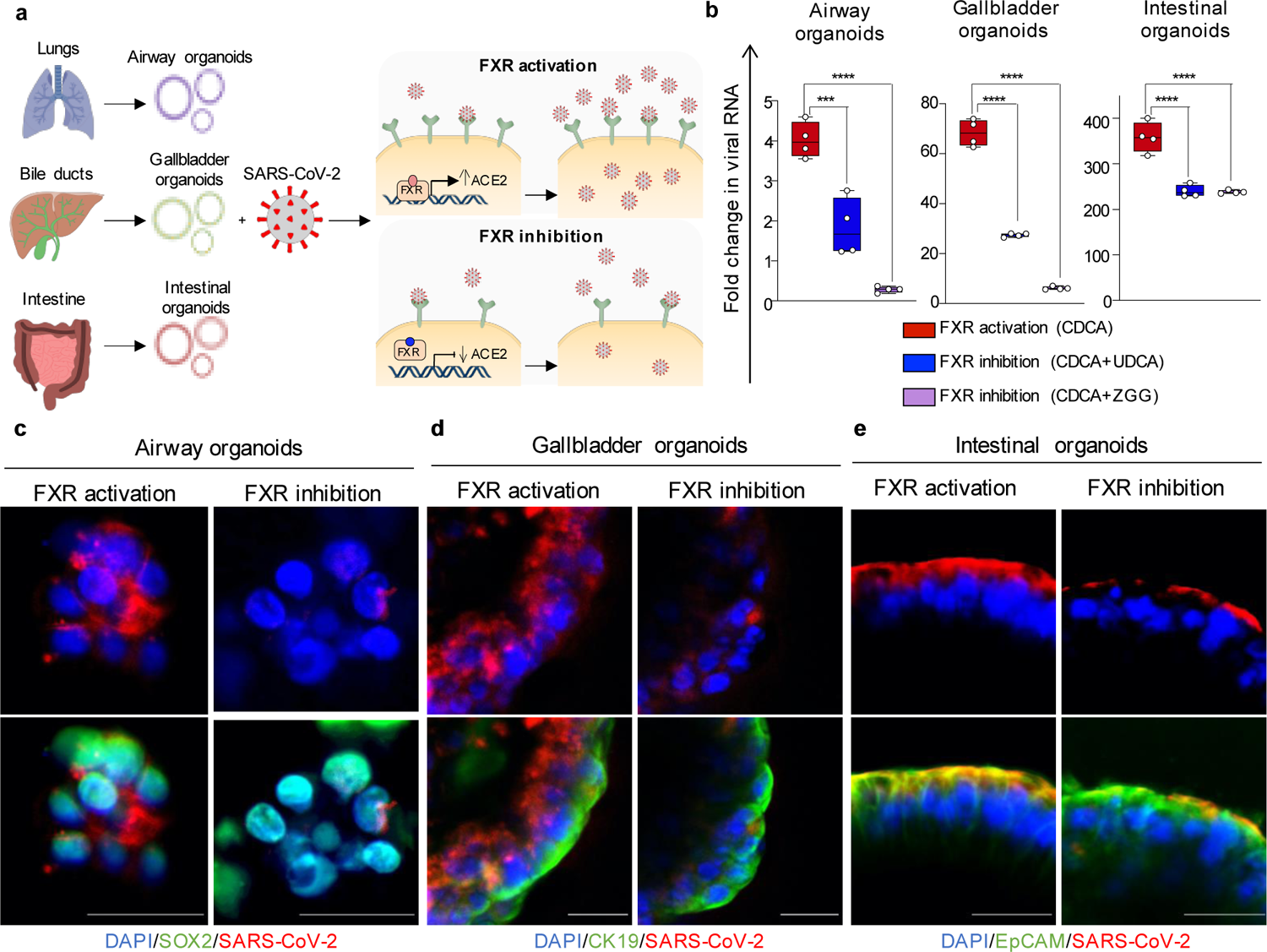
FXR inhibition reduces SARS-CoV-2 infection *in vitro*. (**a**) Schematic illustration of the effect of FXR inhibition on SARS-CoV-2 infection. (**b**) QPCR quantifying fold change in SARS-CoV-2 viral RNA 24 hours post infection (hpi) in primary organoids from different COVID-19 affected tissues. The organoids were treated with physiological levels of bile acids (CDCA) in the presence or absence of FXR inhibitors (UDCA/ZGG). Housekeeping gene, GAPDH; n=4 independent experiments. One-way ANOVA with Dunnett’s correction for multiple comparisons; ***, *P*<0.001; ****, *P*<0.0001; center line, median; box, interquartile range (IQR); whiskers, range; error bars, standard deviation. (**c-e**) Immunofluorescence for the SARS-CoV-2 spike protein 24 hpi in organoids from different COVID-19 affected tissues corresponding to (**b**). Scale bars 25 μm.

### UDCA limits SARS-CoV-2 infection in a human organ *ex vivo*

We then looked to validate these observations using a more physiological setting. In patients, FXR inhibitors are metabolized by the liver, and ultimately distributed to different tissues through systemic circulation. To simulate this process, we performed *ex situ* normothermic perfusion (ESNP) on a human liver graft not used for transplantation (Fig. 3a-b) and characterized the impact of UDCA administered in the circulating blood on ACE2 expression and viral infection at whole organ level. ESNP was developed to enhance organ preservation and reduce reperfusion injury by perfusing organs with warm oxygenated blood prior to transplantation^31^. This setting ensured that the organs used were maintained in near physiological conditions during the experiment^32^. To assess the effect of UDCA, we first measured baseline ACE2 activity in the circulating blood (n=4 independent measurements; Fig. 3c). Baseline ACE2 tissue expression was also assessed using QPCR and immunofluorescence (n=4 independent samples; Fig 3d-e). We then administered UDCA ‘systemically’ in the circulating blood (perfusate). UDCA was diluted to 2000 ng/ml corresponding to the steady-state plasma concentration achieved in patients after multiple doses of oral UDCA^33^. We continued *ex-situ* perfusion with UDCA for 12 hours and obtained repeat blood and tissue samples, matching our pre-UDCA measurements. We observed that *ex vivo* treatment with UDCA reduced ACE2 expression in gallbladder cholangiocytes and ACE2 activity in the serum, compared to the start of the experiment (Fig. 3c-e). To ensure that the observed differences could be attributed to UDCA, we repeated this experiment in a second graft without UDCA treatment. ESNP in the absence of UDCA did not significantly alter these parameters (Extended Data Fig. 8a-c). These results show that clinical doses of circulating UDCA can decrease ACE2 levels in human tissue and serum *ex vivo*.

**Figure 3.**
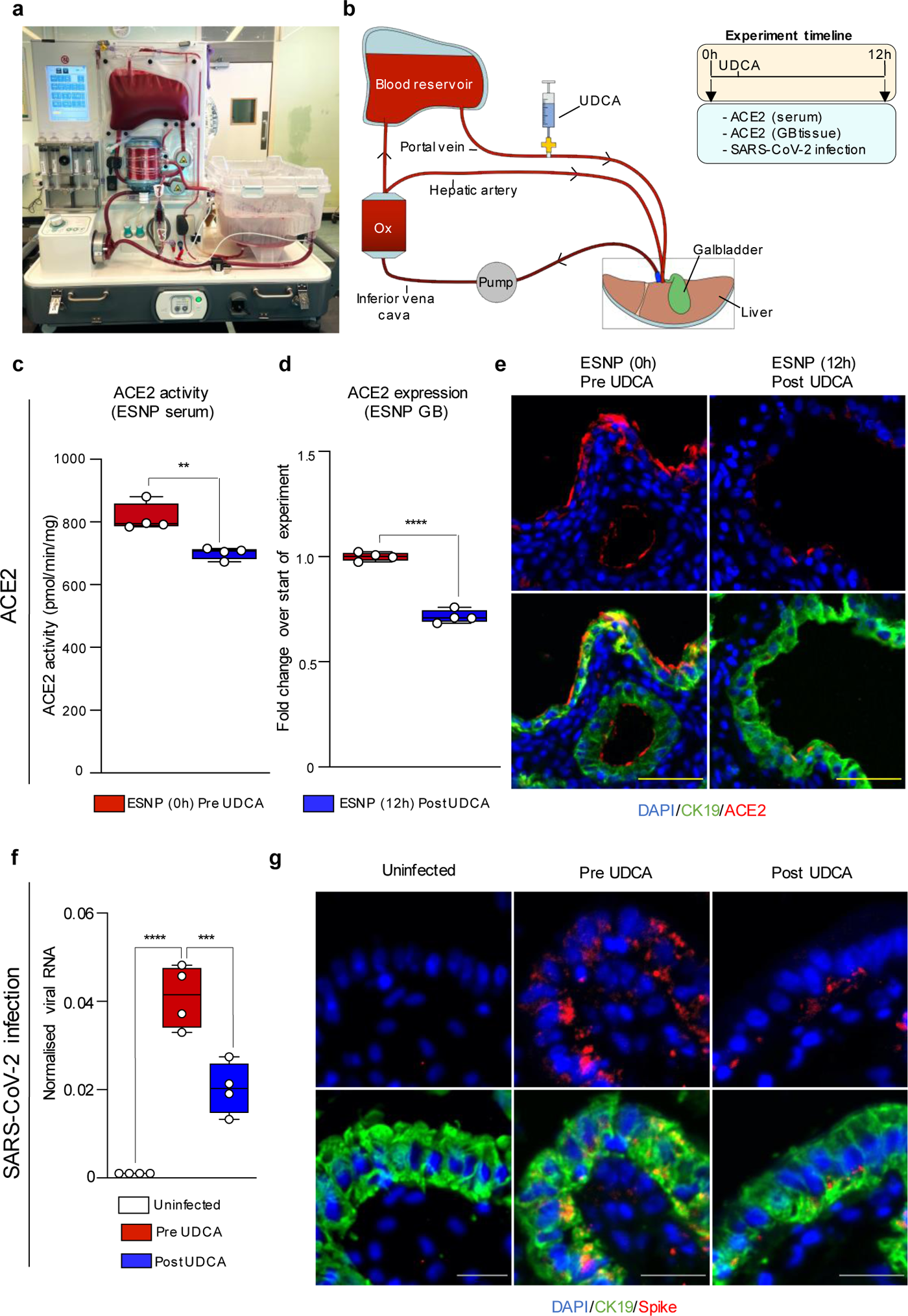
FXR inhibition reduces SARS-CoV-2 infection in a human organ *ex vivo*. (a-**b**) Photograph (**a**) and schematic representation (**b**) of the *ex-situ* normothermic perfusion (ESNP) circuit employed and the experimental timeline. (**c-e**) ACE2 enzymatic activity (**c**), QPCR (**d**) and immunofluorescence (**e**) demonstrating that UDCA treatment reduces ACE2 levels in ESNP. These data correspond to Extended Data Fig. 8a-c showing that ESNP does not modify ACE2 levels in organs not receiving UDCA treatment used as controls. Housekeeping gene, HMBS; n=4 samples. Unpaired two-tailed t-test; **, *P*<0.005; ****, *P*<0.0001; center line, median; box, interquartile range (IQR); whiskers, range; error bars, standard deviation. Scale bars 50 μm. (**f-g**) QPCR (**f**) and immunofluorescence (**g**) showing that UDCA reduces SARS-CoV-2 infection in human gallbladder *ex vivo.* Housekeeping gene, GAPDH; n=4 samples. Scale bars 25 μm. One-way ANOVA with Dunnett’s correction for multiple comparisons; ***, *P*<0.001; ****, *P*<0.0001; center line, median; box, interquartile range (IQR); whiskers, range; error bars, standard deviation.

We then assessed the importance of this reduction in ACE2 for SARS-CoV-2 infection. We infected four independent samples from the graft’s gallbladder before UDCA administration vs. four corresponding independent samples after 12 hours of ESNP treatment with UDCA and observed that UDCA treatment reduced SARS-CoV-2 infection (Fig. 3f-g). These results confirm that FXR inhibition via systemic administration of the approved drug UDCA can downregulate ACE2 in machine-perfused organs and reduce SARS-CoV-2 infection *ex vivo*.

### Clinical observations suggest a correlation between UDCA administration, ACE2 expression and COVID-19 outcome

Our previous results encouraged us to assess the potential impact of UDCA on ACE2 levels in humans. For this, we took advantage of the drug’s extensive use for cholestatic liver disorders (e.g., first-line treatment for the cholestatic autoimmune disorder primary biliary cholangitis, PBC) (Fig. 4a), which allowed us to access serum and tissue samples from patients already receiving UDCA. We first compared ACE2 levels in the gallbladder and common bile duct from PBC liver explants vs. patients with other liver disease not receiving UDCA (n=4 per group) (Fig. 4a) and observed that UDCA treatment was associated with lower levels of ACE2 (Fig. 4b-d). To reinforce this, we measured serum ACE2 activity in patients commencing UDCA for independent clinical reasons before and after administration of UDCA (Fig. 4e; Supplementary Table S4). UDCA was given at the standard therapeutic dosage of 15 mg/kg split over 3 equal doses per day^30^. Serum ACE2 activity in three individuals not receiving UDCA was measured as a control, to account for variance in ACE2 levels over time. We observed reduced ACE2 activity in response to UDCA treatment, while ACE2 activity in control samples remained unchanged (Fig. 4f). Furthermore, we observed a similar reduction in ACE2 activity when comparing unmatched serum samples between UDCA naïve PBC patients (n=11) vs. PBC patients established on UDCA (n=6) (Fig. 4g-h). These experiments suggest that UDCA treatment may reduce ACE2 levels in human (Fig. 4b-h). However, given the potential for confounding bias in any study which is not a clinical trial, it is crucial that these results are validated in the future in a large prospective clinical trial.

**Fig. 4.**
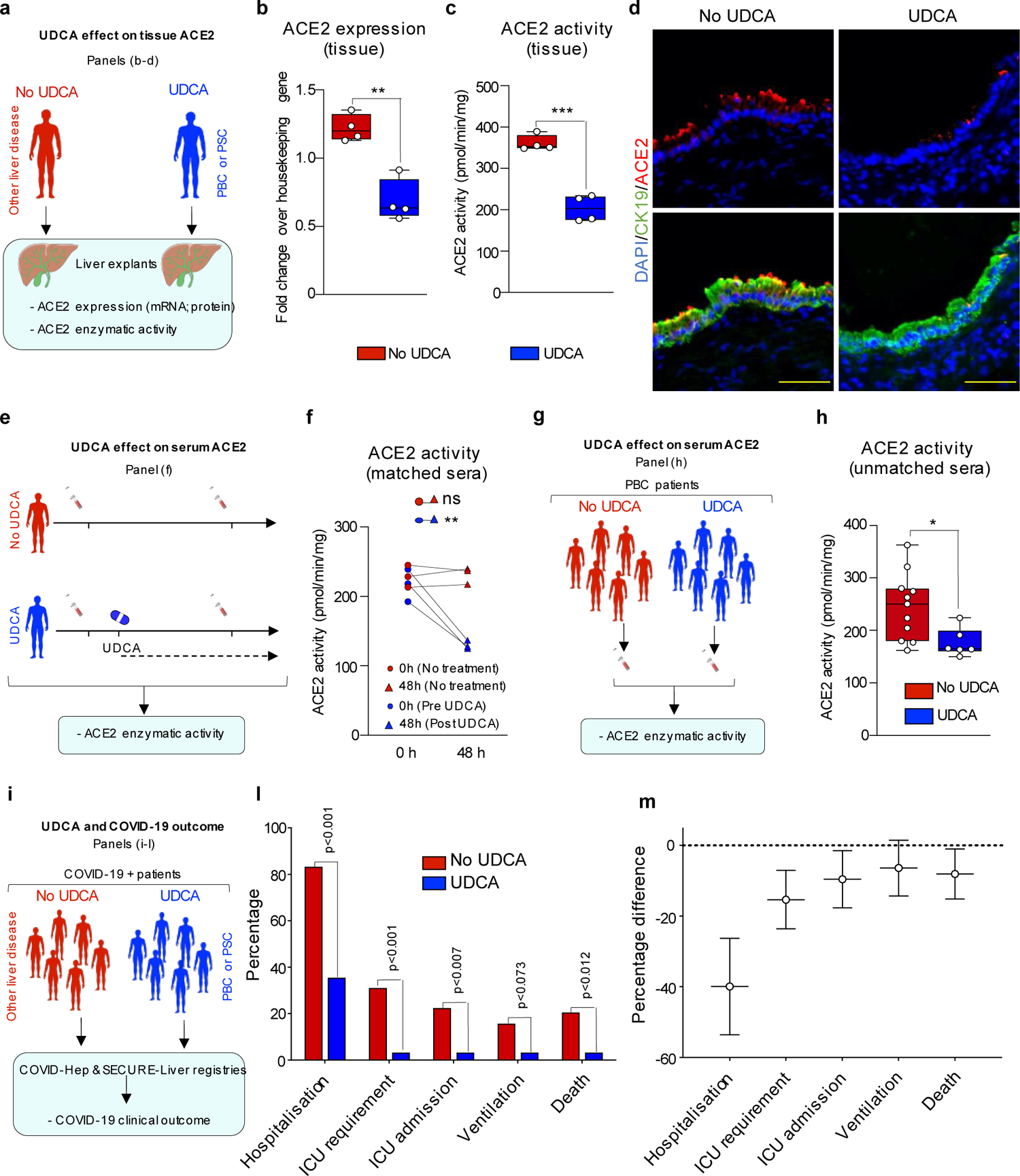
UDCA is associated with lower ACE2 levels and better clinical outcome in COVID-19 patients. (**a**) Schematic illustrating the experimental design corresponding to **(b-d)**. (**b-d**) QPCR (**b**), ACE2 enzymatic activity (**c**) and immunofluorescence (**d**) demonstrating that UDCA treatment is associated with reduced ACE2 levels and activity in tissue from patient treated with UDCA. Housekeeping gene, HMBS; n=4 independent samples. Unpaired two-tailed t-test; *\**\*, *P*<0.005; ***, *P*<0.0001; center line, median; box, interquartile range (IQR); whiskers, range; error bars, standard deviation. Scale bars 50 μm. (**e**) Schematic illustrating the experimental design corresponding to (**f**). (**f**) ACE2 enzymatic activity measurements in patient serum before and after commencing treatment with UDCA (n=3 patients) vs. control patients not receiving UDCA (n=3 patients). UDCA treatment was started for independent clinical indications. Paired two-tailed t-test; ns, non-significant; **, *P*<0.005. Lines connect matched serum samples from the same individual at 0h and 48h. (**g**) Schematic illustrating the experimental design corresponding to (**h**). (**h**) ACE2 enzymatic activity in unmatched serum samples from UDCA naïve PBC patients (n=11 patients) vs. PBC patients established on UDCA (n=6 patients). Unpaired two-tailed t-test; *, *P*<0.05; center line, median; box, interquartile range (IQR); whiskers, range; error bars, standard deviation. These data correspond to Supplementary Table S4. (**i**) Schematic illustrating the experimental design corresponding to (**l**-**m**). (**l**) Clinical outcome from COVID-19 in patients with chronic liver disease in the presence or absence of UDCA treatment. (**m**) Propensity-score matched analyses showing major outcomes following SARS-CoV-2 infection in patients taking UDCA compared to non-UDCA controls. The data correspond to Supplementary Table S5 (patient characteristics).

Based on these observations, we decided to explore the potential impact of UDCA treatment on the outcome of COVID-19 in patients. To explore this, we interrogated the COVID-Hep/SECURE-Liver registries^34–36^ which comprise data of patients with chronic liver disease (n=1,096) and who developed COVID-19, including patients with cholestatic liver disorders receiving UDCA (n=31) (Fig. 4i) (Supplementary Table S5). We observed that, accepting the potential for selection bias in case reporting, patients receiving UDCA had better outcomes compared to patients not receiving UDCA, including reduced hospitalisation, ICU admission, requirements for invasive ventilatory support and death (Fig. 4l), including after propensity score matching for sex, age and stage of liver disease (Fig. 4m). These results suggest that UDCA may be associated with better clinical outcomes of COVID-19 and could warrant the development of future clinical trials.

## Discussion

Considered collectively, our findings demonstrate that FXR participates in the regulation of ACE2 expression in multiple tissues involved in COVID-19. FXR inhibition, using the clinically approved drug UDCA, downregulates ACE2 expression and reduces SARS-CoV-2 infection *in vitro* and *ex vivo*. Furthermore, our clinical observations raise the possibility that UDCA may reduce ACE2 levels in patient serum and tissue and suggest a potential correlation between UDCA and positive clinical outcomes in COVID-19 patients.

The finding that FXR regulates ACE2 is novel but not entirely surprising. The functions of ACE2 as a molecular chaperone for the amino acids transporter SLC6A19^37^ and as a peptidase justify its presence in the GI tract and suggest a potential role in digestion. Accordingly, the upregulation in ACE2 expression by FXR, which is activated by bile, a digestive fluid, may reflect a mechanism to increase peptidase levels and amino acids absorption during digestion. Furthermore, in addition to its role in the GI system, FXR is expressed in multiple organs, including the lungs^24, 38^, with a broad variety of functions^23, 39–44^, while its natural ligands, such as bile acids and hormones^45^ (e.g., androgen^46^) are present in the systemic circulation^26^. This broad expression explains how FXR could regulate ACE2 in multiple tissues beyond the biliary tree.

Our results illustrate the potential of ACE2 modulation as a novel host-directed therapy against COVID-19. Because host-directed treatments do not target the virus, they are less susceptible to resistance and more likely to be effective against multiple viral variants^47^.

Furthermore, ACE2 is a common receptor for multiple coronaviruses, such as SARS-CoV and HCoV-NL63. Thus, ACE2 modulation might prove to be an effective therapeutic approach for future pandemics. Finally, large Mendelian randomization analyses in over 7,554 patients hospitalised with COVID-19 and more than 1 million controls demonstrated that higher ACE2 levels strongly correlated with increased risk of COVID-19 hospitalisation. These data support our observations on the effects of ACE2 modulation for SARS-CoV-2 infection and identify ACE2 modulators as priority candidates for evaluation in COVID-19 trials^11^.

Our finding that FXR inhibition through UDCA or ZGG reduces ACE2 expression, and limits SARS-CoV-2 infection, identifies a new potential clinical application for FXR inhibitors. UDCA is easy to administer orally, well-tolerated, with a favourable safety profile, easily stored, affordable and accessible to health systems world-wide for large scale production, as it is off patent. Our results also show that UDCA reduces SARS-CoV-2 infection *in vitro* and *ex vivo* and our clinical observations suggest that it could be associated with lower ACE2 and improved outcomes in COVID-19. Nevertheless, our study is not a clinical trial and therefore we cannot exclude the potential for confounding and selection biases. Consequently, it will be imperative to validate these results in prospective double blinded clinical trials and fully assess the impact of this drug on ACE2 levels and SARS-CoV-2 infection.

Finally, we demonstrated that UDCA could reduce ACE2 levels and SARS-CoV-2 infection in machine perfused organs. This is one of the first studies testing the effect of a drug in a whole human organ perfused *ex situ*. This finding could prove important for organ transplantation, especially given concerns about peri-operative viral transmission^48^. Furthermore, although more data are required to definitively establish this approach, our work sets the stage for future studies using machine-perfused organs for pharmacological studies.

In conclusion, these results validate CDCA-treated cholangiocytes organoids as a novel platform for disease modelling and drug testing against SARS-CoV-2 infection; identify FXR as a new therapeutic target in the management of COVID-19 and open up new avenues for the modulation of ACE2 using FXR inhibitors for prevention and therapy of SARS-CoV-2 infection.

## Supporting information

Brevini et al., Supplementary Information

## Methods

### Ethical approval

All human samples were obtained from patients, deceased transplant organ donors or liver explants with informed consent for use in research, and ethical approval (Research Ethics Committee - REC 09/H0305/68; 14/NW/1146; 15/EE/0152; 15/WA/0131; 18/EE/0269; and Papworth Hospital Research Tissue Bank project number T02233). Human livers retrieved for transplantation but subsequently declined were used for *ex situ* normothermic perfusion experiments (National Research Ethics Committee East of England – Cambridge East 14/EE/0137). The COVID-Hep.net and SECURE-Liver registries data were deemed not to constitute human research by Clinical Trials and Research Governance at the University of Oxford (https://covid-hep.net/img/CTRG_COVID-Hep_20200402.pdf) and by the Institutional Review Board of University of North Carolina (https://covidcirrhosis.web.unc.edu/faq/) respectively.

### 10x single cell RNA sequencing, data analysis and availability

We used our previously published single cell RNA sequencing (scRNAseq) dataset including primary cholangiocytes, cholangiocyte organoids originating from different regions of the biliary tree (intrahepatic ducts, common bile duct and gallbladder), and the same organoids following bile treatment. Tissue dissociation, cell isolation, 10X single cell library preparation and 10X data processing, normalisation and analysis was performed as previously described^18^. 10X raw data (fastq files) have been deposited in the repository ArrayExpress with the accession number E-MTAB-8495.

### Human tissue collection and processing

Human primary tissue was obtained from biopsies, deceased transplant organ donors or liver explants after obtaining informed consent. Depending on the application, primary fresh tissue was embedded in OCT (Optimal Cutting Temperature) compound and stored at −80°C; or fixed in 10% formalin, dehydrated and embedded in paraffin. Sections from embedded tissue were cut at a thickness of 5-10 μm using a cryostat or a microtome and mounted on microscopy slides for further analysis.

### Bile sample collection and processing

Human bile was collected during ERCP (Endoscopic Retrograde Cholangio-Pancreatography) or intraoperatively with informed consent from the patient. For viral RNA quantification samples were immediately lysed using an equal volume of RNA lysis buffer (Sigma) and stored at −20°C.

### Cell culture

Primary cholangiocytes were isolated and cholangiocyte organoids were derived and cultured using our established methodology^18, 49, 50^. Cholangiocyte organoids obtained from intrahepatic ducts (IHD), common bile duct (CBD) and gallbladder (GB) tissue were used in this study. Cholangiocytes derived from any of the different regions of the biliary tree (IHD, CBD, GB) acquired a common gallbladder identity when treated with CDCA, as previously reported^18^. The experiments described were performed with cholangiocyte organoids derived from all the three regions of the biliary tree (IHD, CBD and GB) and provided congruent results. For consistency, the results shown correspond only to cholangiocyte organoids derived from the gallbladder (gallbladder cholangiocyte organoids – GCOs).

Human primary intestinal organoids, derived from terminal ileum biopsies were provided by Simon Buczacki’s group. The organoids were derived following a modification of previously described protocols^51^, embedded in Matrigel and cultured in Intesticult (StemCell Technologies) supplemented with Penicillin-Streptomycin and Rho kinase inhibitor (Stratech Scientific). Human primary airway organoids were provided by Joo-Hyeon Lee’s lab. The organoids were derived and cultured as previously described^28^. Vero E6 cells (ATCC™ CRL – 1586) were grown on tissue culture plates or T25 flasks in 10% FBS DMEM supplemented with L-glutamine and Penicillin-streptomycin as previously described^52^.

### Biological materials availability

Detailed protocols for the derivation of primary organoids have been previously reported^28, 50^. Cell lines are available from standard commercial sources (https://www.lgcstandards-atcc.org - Vero E6 cells, ATCC™ CRL – 1586).

### Modulation of FXR activity

Chenodeoxycholic acid (CDCA) and ursodeoxycholic acid (UDCA) were purchased from Sigma Aldrich (C9377-5G and U-5127-5G), while Z-Guggulsterone (ZGG) was purchased from Santa Cruz (sc-204414) and reconstituted following the manufacturer’s instructions. To modulate FXR activity, organoids were incubated with a final concentration of 10 μM CDCA; or 10 μM CDCA in combination with 10 μM of UDCA or ZGG.

### Chromatin immunoprecipitation

Approximately 6 × 10^5^ cells were used for each chromatin immunoprecipitation (ChIP), and cells were incubated with fresh medium with 100 μM of CDCA, UDCA or ZGG 2 h before collection. ChIP was performed using the True Micro ChiP kit (Diagenode C01010130) according to manufacturer’s instructions. In brief, following pre-clearing, the lysate was incubated overnight with the FXR antibody (Santa Cruz sc-25309 X) (Supplementary Table S1) or non-immune IgG. ChIP was completed and immunoprecipitated DNA was purified using MicroChip DiaPure columns (Diagenode C03040001). Samples were analysed by QPCR using the ΔΔ*C*t approach as previously described^50^ (see Supplementary Table S3 for primer sequences). Primers flanking the FXR responsive element (FXR RE) on the well-known FXR target gene *OST*α^53^ were used as positive control, while primers flanking a site distant from the FXR RE on the ACE2 promoter were used and a negative control. The results were normalized to the enrichment observed with non-immune IgG ChIP controls.

### Immunofluorescence, RNA extraction and QPCR

Immunofluorescence, RNA extraction and quantitative real-time PCR (QPCR) were performed as previously described^49, 50, 54, 55^. A complete list of the primary and secondary antibodies used is provided in Supplementary Table S1. A complete list of the primers used is provided in Supplementary Table S2.

All QPCR data are presented as the median, interquartile range (IQR) and range (minimum to maximum) of four independent experiments unless otherwise stated. Values are relative to the housekeeping gene Hydroxymethylbilane Synthase (*HMBS*) or Glyceraldehyde-3-Phosphate Dehydrogenase (*GAPDH)*. Statistical analysis is described in the relevant section. For comparative immunofluorescence images, the cells or sections being compared were stained simultaneously, using the same primary and secondary antibody master mix. All immunofluorescence images were acquired using a Zeiss LSM 700 confocal microscope.

The same laser power and exposure settings were used to acquire comparative images. Imagej 2.0.0 software (Wayne Rasband, NIHR, USA, http://imagej.nih.gov/ij) was used for image processing. Each immunofluorescence image is representative of at least 3 different experiments.

### Flow cytometry analyses

Flow cytometry in organoids was performed as previously described^50^. In summary, organoids were collected using Cell Recovery Solution (Corning) for 20 min at 4 °C and were then centrifuged at 444*g* for 4 min and dissociated to single cells using StemPro Accutase (Invitrogen). Cells were subsequently fixed using 4% paraformaldehyde (PFA) for 20 min at 4 °C. All flow cytometric analyses were performed on a BD LSR-II flow cytometer from BD Biosciences and analysed using FlowJo v.10.4.2. The gating strategy is provided in Supplementary Figure S1.

### SARS-CoV-2 isolate

The SARS-CoV-2 virus used in this study is the clinical isolate named “SARS-CoV-2/human/Liverpool/REMRQ0001/2020”^52^ derived from a patient’s naso-pharyngeal swab and isolated by Lance Turtle (University of Liverpool) and David Matthews and Andrew Davidson (University of Bristol).

### SARS-CoV-2 infection

All work with infectious SARS-CoV-2 was performed under containment level 3 (CL-3) conditions at the Cambridge institute of Therapeutic Immunology and Infectious Diseases (CITIID). SARS-CoV-2 was gifted to the users of the CITTID CL-3 by Ian Goodfellow^56, 57^ and propagated on Vero E6 cells as previously described^52^. Viral titration was determined using the TCID50 method on Vero E6 cells^52^. For viral infection primary organoids were passaged and incubated with SARS-CoV-2 in suspension at a multiplicity of infection (MOI) of 1 for 2 hours. Subsequently, the infected organoids were washed twice with 10 ml of culture media to remove the viral particles. Washed organoids were plated in 40 μl Matrigel domes, cultured in organoid medium and harvested at different timepoints.

To test whether SARS-CoV-2 produced by infected COs retained its infective capacity, the supernatant from infected COs was collected at 24 hours post infection and used to infect a fresh batch of SARS-CoV-2 naive organoids.

### Fixation of SARS-CoV-2 infected organoids/tissue

Organoids for IF were cultured on coverslips, placed at the bottom of the wells of a 24-well plate. The culture medium was aspirated and replaced with 500 µl of 8% PFA for a minimum of 30 minutes. Following fixation, the coverslips were recovered, transferred to a clean plate, and fresh PBS was added. Primary tissue was fixed for a minimum of 4 hours with 8% PFA and then transferred to a clean plate with fresh PBS.

### Quantification of viral infection

Organoids or primary tissue were infected in 24-well plates as described above. Total RNA samples were prepared by adding 500 µl of lysis buffer (25 mM Tris-HCL+ 4 M Guanidine thiocyanate with 0.5% β-mercaptoethanol) to each well and transferring the lysate (1 ml) to a 5 ml Eppendorf tube. Tubes were vortexed, and 100% analytical grade ethanol was added to a final concentration of 50%. After 10 minutes of incubation, 860 µl of lysis buffer (containing MS2 bacteriophage as an internal extraction and amplification control) were added and thoroughly mixed. The RNA was then isolated using an RNA spin column as previously described^58^. Viral replication was quantified using QPCR for the expression of the viral RNA-dependent RNA polymerase (*RdRp*) gene with primers specific for a 222 bp long fragment from a conserved region of the gene. *GAPDH* was used as a housekeeping gene and MS2 was used as an internal reference as previously described^58^. Viral load was determined relative to *GAPDH*. The fold change over the starting viral RNA was measured to quantify increases in viral RNA indicative of viral replication. The sequences of primers/probes used are provided in Supplementary Table S2.

### Transmission Electron Microscopy

Infected organoids were fixed in 4% paraformaldehyde; 2.5% glutaraldehyde in 0.1M sodium cacodylate buffer overnight at 4°C, washed and stored in 0.1M sodium cacodylate buffer before processing. Samples were postfixed in 1% aqueous osmium tetroxide (TAAB, UK); 1.5% potassium ferricyanide overnight at 4°C, washed thoroughly in dH_2_O and *en bloc* stained in 3% aqueous uranyl acetate (Agar Scientific, UK) for 24h at 4°C. Samples were dehydrated through an ethanol series, infiltrated with 1:1 propylene oxide:resin (TAAB, UK) and blocks of fresh resin polymerised at 60°C for 48h. Ultrathin sections of ∼60nm were cut from blocks using an EM UC7 ultramicrotome (Leica Microsystems, UK) and mounted on copper grids coated with carbon and formvar (Agar Scientific, UK). Grids were post-stained in unrayl acetate and lead citrate and imaged using a HT7800 transmission electron microscope (Hitachi High Technologies, Japan) operating at 100kV.

### *Ex situ* normothermic perfusion (ESNP) of donor livers

The OrganOx metra normothermic liver perfusion device was used for *ex situ* perfusion of human livers as previously described^18, 31, 32^. The machine, which is clinically used for preservation of livers for transplantation enables prolonged automated organ preservation by perfusing it with ABO-blood group-matched normothermic oxygenated blood. The perfusion device incorporates online blood gas measurement, as well as software-controlled algorithms to maintain pH, PO2 and PCO2 (within physiological limits), temperature, mean arterial pressure and inferior vena cava pressure within physiological normal limits.

Donor livers not used for transplantation were maintained with ESNP and treated with UDCA for 12 hours. In brief, the hepatic artery, portal vein, inferior vena cava and bile duct were cannulated, connected to the device and perfusion commenced. UDCA was resuspended in 0.9% (w/v) NaCl solution and injected in the blood circuit to achieve a final concentration of 2000 ng/ml, which is the steady state concentration of UDCA detected in serum after multiple doses of UDCA^33^. Samples were collected before and 12 hours after UDCA administration as described in the results section ‘UDCA limits SARS-CoV-2 infection in a human organ *ex vivo*’.

### *Ex vivo* SARS-CoV-2 infection of human gallbladder tissue

Freshly obtained gallbladder tissue was processed into small rectangular pieces of 0.5×0.5 cm and were rinsed withWilliam’s E medium with supplements as previously described^50^. Washed specimens were placed in wells of a 24-well plate (one specimen per well) and infected with SARS-CoV-2. An inoculum of 1.2×10^5^ PFU/ml at 500 µl per well was used. After two hours, the inoculum was removed, and the specimens were washed three times with PBS. The infected human gallbladder tissue was then cultured in 500 µl of William’s E medium with supplements. Supernatant and tissue were harvested for QPCR and immunofluorescence at 2 and 24 hours post infection.

### Blood sample collection and processing

Blood samples were collected from patients as part of the UK-PBC Nested Cohort study after obtaining informed consent, anonymised and analysed by a blinded researcher. To obtain serum from full blood, the samples were spun at 4°C at 1000 g for 10 minutes to allow for serum separation and serum was collected as the supernatant.

### ACE2 enzymatic activity

ACE2 enzymatic activity was performed on serum samples and tissue lysates using the ACE2 activity fluorometric kit (abcam ab273297) following manufacturer’s instructions using a SpectraMax M2 (Molecular Devices).

### Patient data

Data for patients with chronic liver disease were collected as described elsewhere^34, 36^. Briefly, collated results from two open online reporting registries (COVID-Hep.net and SECURE-Liver) were examined. Reports were asked to report cases of laboratory confirmed COVID-19 in patients with chronic liver disease at the end of the disease course. Anonymous clinical and demographic data were collected, filtered to remove duplicate entries, those with incomplete records, those with prior liver transplantation, those not over 18 years of age and those without laboratory confirmed infection.

## Statistical analyses

Statistical analyses were performed using GraphPad Prism 9 or Stata 15.1 (StataCorp, College Station, TX, USA). The normal distribution of our values was evaluated using the Shapiro-Wilk test where appropriate. For comparison between two groups, a two-tailed Student’s t-test or the non-parametric Mann-Whitney test were used depending on the normality of our distribution. To compare matched samples from the same individual a two-tailed paired Student’s t-test was used (the normal distribution of our data was confirmed using the Shapiro-Wilk test). Variance between samples was tested using the Brown-Forsythe test. For comparing multiple groups to a reference group one-way ANOVA followed by Dunnett’s test was used between groups with equal variance. Immunofluorescence images are representative of 4 independent experiments. Data are represented in box plots and elements are defined as follow: center line, median; box, interquartile range (IQR); whiskers, range; error bars, standard deviation.

For registry data, comparisons between proportions were calculated using two-tailed Fisher’s exact tests in a method similar to that reported elsewhere^34, 36^. For propensity score-matched analyses, 1:10 matched samples (using the nearest neighbour approach) were constructed with hospitalisation, physician-reported requirement for intensive care, intensive care admission, mechanical ventilation, and death as the outcome variables. Covariables used were age, sex and categorical stage of chronic liver disease according to the *Child-Turcotte-Pugh* class. Propensity score matching was performed using the *teffects* function in Stata. The average treatment effect on the treated (ATET) was calculated with robust Abadie-Imbens standard errors and derived 95% confidence intervals are presented.

## Data and materials availability

All data are available in the main text or the Extended Data. Single-cell RNA sequencing data are available on ArrayExpress. Accession number: E-MTAB-8495.

## Acknowledgements

The authors would like to thank the European Association for the Study of the Liver (EASL) and the American Association for the Study of Liver Disease (AASLD) for supporting the COVID-Hep and SECURE-Liver registries; Prof Stefan Marciniak and Prof Paul J. Lehner for comments and feedback on the manuscript; Prof Ian Goodfellow for providing the viral isolate; Dr. Mark Wills and Dr. Simon Clare for all their work ensuring a safe CL3 working environment; Claire Cormie and Stephanie Brown for general lab support; Peter Humphreys, Darran Clement and the Jeffrey Cheah Biomedical Centre Imaging Facility for their help with confocal and transmission electron microscopy; the NIHR Cambridge BRC Cell Phenotyping Hub for their help with flow cytometry and processing of samples; the building staff of the Jeffrey Cheah Biomedical Centre for maintaining the institute open and safe during the period of lockdown; Janeane Hails, Konstantina-Irene Nikitopoulou and Abigail Ford for collecting blood samples; Maria Colzani for advising on flow cytometry; Anne Wiblin from Abcam for advising on antibodies; the Cambridge Biorepository for Translational Medicine for the provision of human tissue used in the study.

## Author contributions

T.B. conceived and designed the study, performed experiments, acquired, interpreted and analysed the data, developed and validated the protocols described, generated the figures, wrote and edited the manuscript. M.M. performed SARS-CoV-2 infection experiments, collected and processed infectious samples, contributed to data interpretation and critical revision of the manuscript for important intellectual content. G.J.W. acquired, analysed and interpreted patients’ data from the COVID-Hep and SECURE-Liver registries and contributed to critical revision of the manuscript for important intellectual content. W.T.H.G. contributed to sample collection and critical revision of the manuscript for important intellectual content. S.F. and P.M. contributed to viral infection experiments. S.D. performed transmission electron microscopy sample processing and preparation. S.V., M.D.D. and T.W.M.C. expanded and provided cell lines, contributing to cell culture. V.L.M. contributed to blood sample collection. R.E.K. and T.L.W. provided primary tissue samples. V. G., M.V.G and O.C.T. provided samples for QPCR. D.M. performed bioinformatic analyses. J.B. provided primary tissue. S.S. and S.S.U. contributed to critical revision of the manuscript for important intellectual content. L.S. provided perfusionist support for ESNP experiments. K.S.-P. provided primary tissue and established livers on ESNP. S.E.D. provided primary tissue and contributed to histological analyses. T.M., E.B., A.W.L., A.M.M. and A.S.B.IV acquired and analyzed patient data included in the COVID-Hep and SECURE-Liver registries. R.K.G. contributed to critical revision of the manuscript for important intellectual content. S.B. and P.L. contributed to critical revision of the manuscript for important intellectual content. G.C. and A.P.D. provided primary tissue samples and contributed to critical revision of the manuscript for important intellectual content. S.J.A.B. and J-H.L. provided primary organoids, contributed to data interpretation and critical revision of the manuscript for important intellectual content. P.G. provided tissue and bile samples and reviewed the manuscript for important intellectual content. A.J.W. and C.J.E.W. provided tissue and bile samples, contributed to ESNP experiments and reviewed the manuscript for important intellectual content. G.F.M. provided samples and contributed to critical revision of the manuscript for important intellectual content and sample provision. G.D. contributed to critical revision of the manuscript for important intellectual content. L.V. designed and conceived the study, interpreted the data and edited the manuscript. F.S. designed and conceived the study, interpreted the data, edited the manuscript and performed ESNP experiments. All the authors approved the manuscript.

## Supplementary Information

Supplementary Information file includes:

**Supplementary Figure S1. Gating strategy for flow cytometry analyses.**

**Supplementary Table S1. List of primary and secondary antibodies.**

**Supplementary Table S2. List of primers used for QPCR.**

**Supplementary Table S3. List of primers used for ChIP-QPCR.**

**Supplementary Table S4. Patient characteristics relative to 4f-h.**

**Supplementary Table S5. COVID-Hep and SECURE-Liver registries patient cohort characteristics.**

---

**Extended Data Figure 1.**
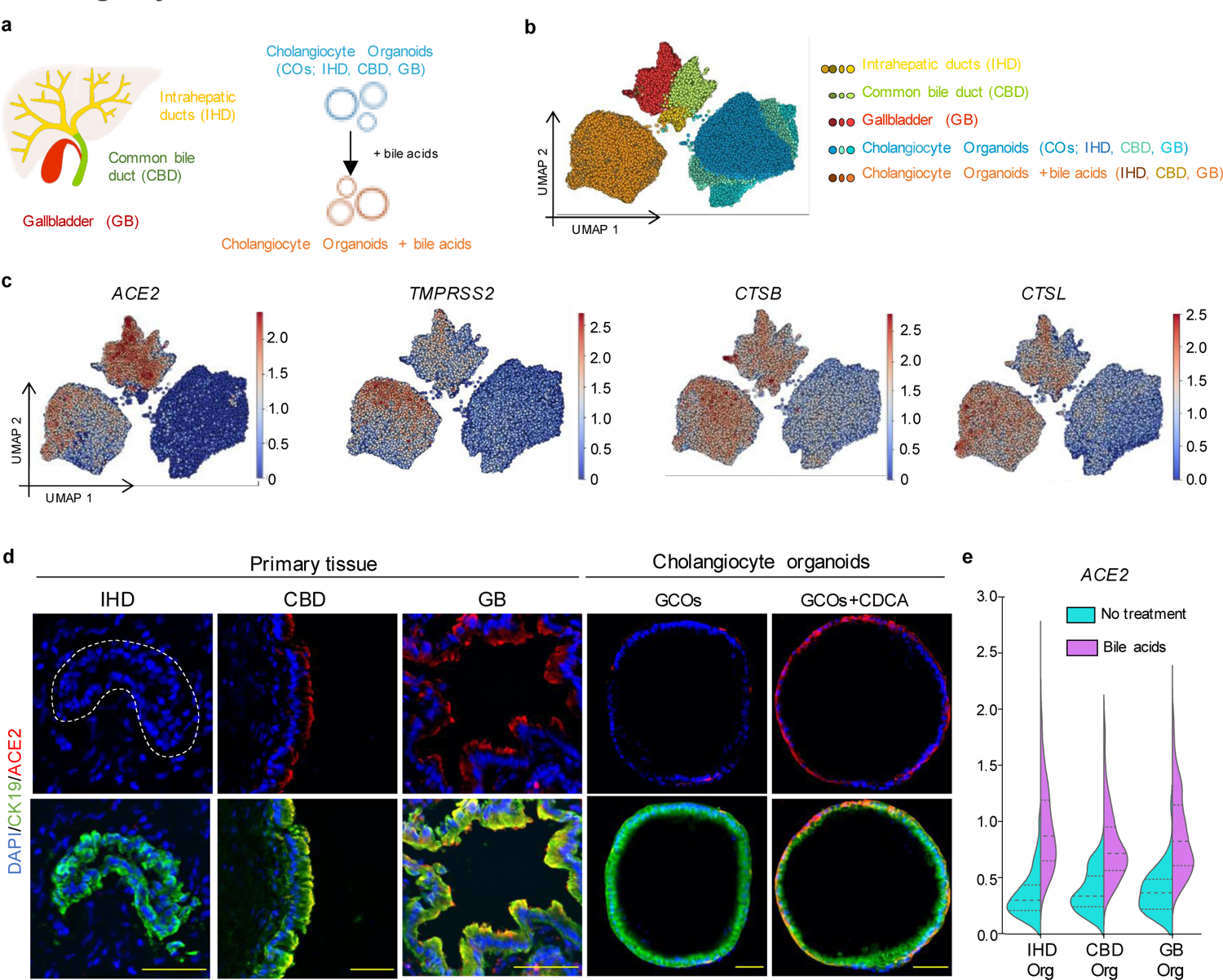
Expression of markers involved in SARS-CoV-2 entry in cholangiocytes. (**a**) Schematic illustration of different cholangiocyte populations corresponding to different areas of the biliary tree and cholangiocyte organoids (COs) derived from different areas of the biliary tree grown in absence or presence of the bile acids. (**b**) UMAP plot illustrating different cholangiocyte populations from (**a**) analysed by scRNAseq. (**c**) UMAP plots showing that viral entry related genes are predominantly expressed in extrahepatic cholangiocytes and COs treated with CDCA. (**d**) Immunofluorescence illustrating that ACE2 is predominantly expressed in extrahepatic cholangiocytes and CDCA-treated gallbladder cholangiocyte organoids (GCOs). Scale bars 50 μm. (**e**) Violin plot showing that cholangiocyte organoids upregulate ACE2 when treated with bile acids regardless of their region of origin.

**Extended Data Figure 2.**
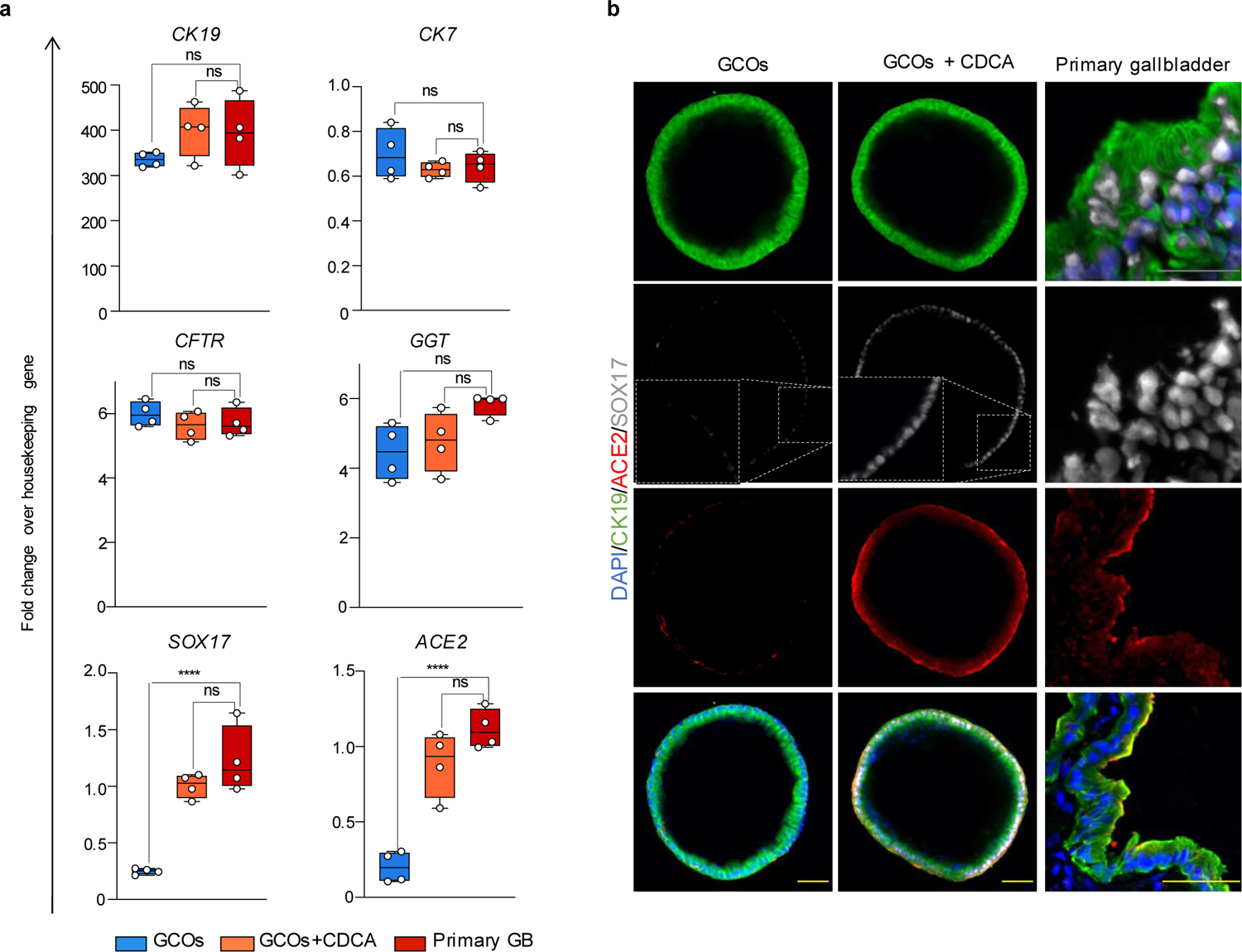
Cholangiocyte organoids treated with CDCA acquire a gallbladder identity and express ACE2. (**a**) QPCR showing that upon treatment with the bile acid CDCA cholangiocyte organoids assume a gallbladder identity, they express the gallbladder marker SOX17 and upregulate the viral receptor ACE2 at levels comparable to primary gallbladder. Housekeeping gene, HMBS; n=4 independent experiments. One-way ANOVA with Dunnett’s correction for multiple comparisons; ns, non-significant; ****, *P*<0.0001; center line, median; box, interquartile range (IQR); whiskers, range; error bars, standard deviation. (**b**) Immunofluorescence images showing that CDCA treatment results in upregulation of the gallbladder marker SOX17 and the viral receptor ACE2 in organoids. Yellow scale bars 50 μm; grey scale bar 25 μm.

**Extended Data Figure 3.**
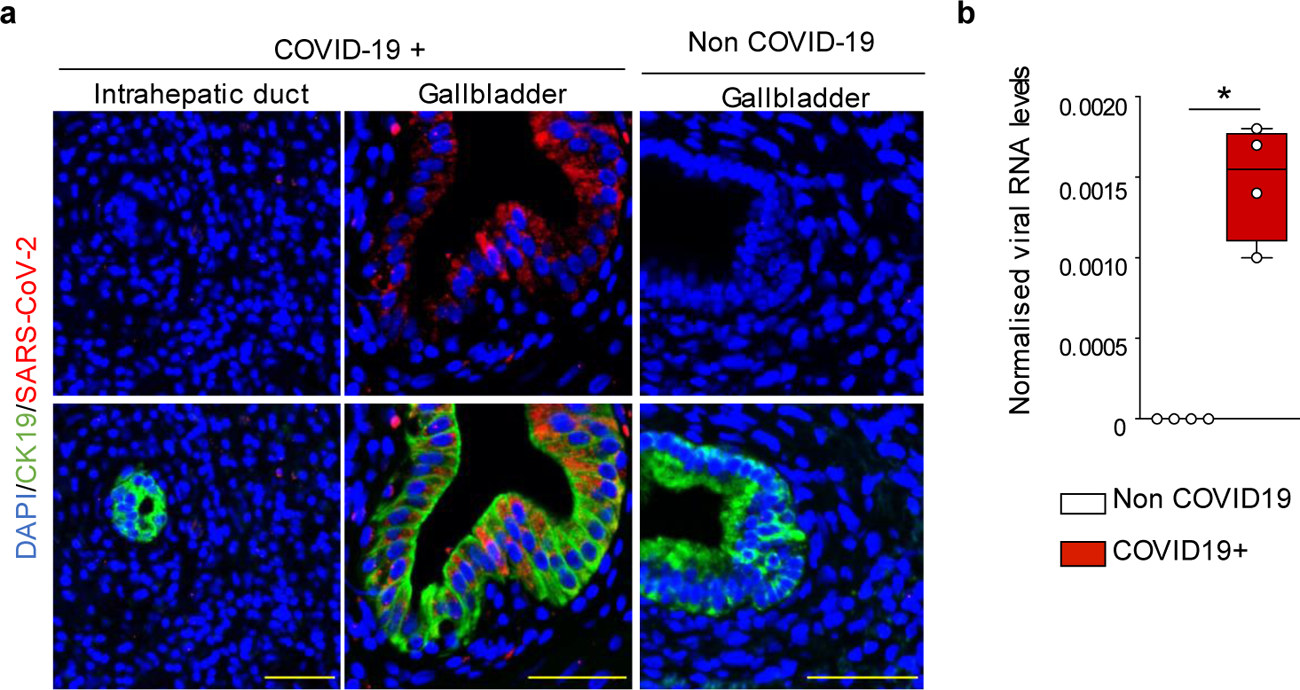
SARS-CoV-2 infects gallbladder cholangiocytes. (**a**) Immunofluorescence illustrating SARS-CoV-2 infection in gallbladder cholangiocytes of COVID-19+ patient but not in intrahepatic cholangiocytes or non-COVID-19 control gallbladder. Scale bars 50 μm. (**b**) QPCR confirming detection of SARS-CoV-2 RNA in bile of COVID-19+ patients. Housekeeping gene, HMBS; n=4 samples. Mann-Whitney test; *, *P*<0.05; center line, median; box, interquartile range (IQR); whiskers, range; error bars, standard deviation.

**Extended Data Figure 4.**
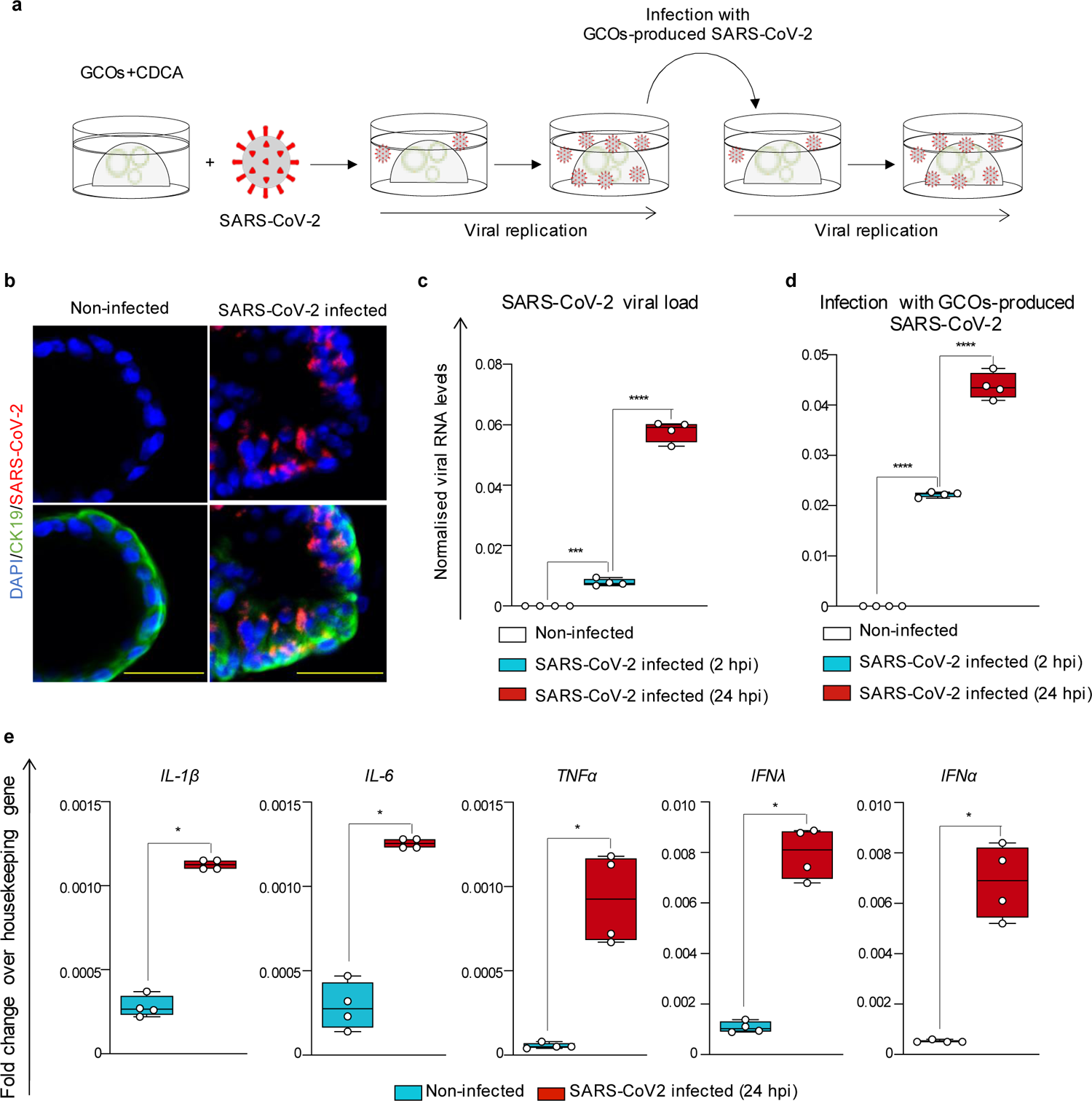
CDCA-treated GCOs can be infected by SARS-CoV-2. (**a**) Schematic illustration outlining the experiments for SARS-CoV-2 infection. (**b**-**c**) Immunofluorescence (**b**) and QPCR (**c**) confirming detection of SARS-CoV-2 in GCOs+CDCA at 24 hours post infection (hpi). Scale bars 50 μm; housekeeping gene, GAPDH; n=4 independent samples. One-way ANOVA with Dunnett’s correction for multiple comparisons; ns, non-significant; ***, *P*<0.005; ****, *P*<0.0001; center line, median; box, interquartile range (IQR); whiskers, range; error bars, standard deviation. (**d**) QPCR showing infection of CDCA-treated GCOs with SARS-CoV-2 harvested from the supernatant of previously infected organoids (24 hpi); illustrating that SARS-CoV-2 produced in GCOs+CDCA retains its infective capacity. Housekeeping gene, GAPDH; n=4 independent samples. One-way ANOVA with Dunnett’s correction for multiple comparisons; ns, non-significant; ***, *P*<0.005; ****, *P*<0.0001; center line, median; box, interquartile range (IQR); whiskers, range; error bars, standard deviation. (**e**) QPCR showing up-regulation of innate immune and anti-viral response genes in GCOs+CDCA following SARS-CoV-2 infection. Housekeeping gene, GAPDH; n=4 independent samples. Two-tailed Mann-Whitney test; ns, non-significant; *, *P*<0.05; center line, median; box, interquartile range (IQR); whiskers, range; error bars, standard deviation.

**Extended Data Figure 5.**
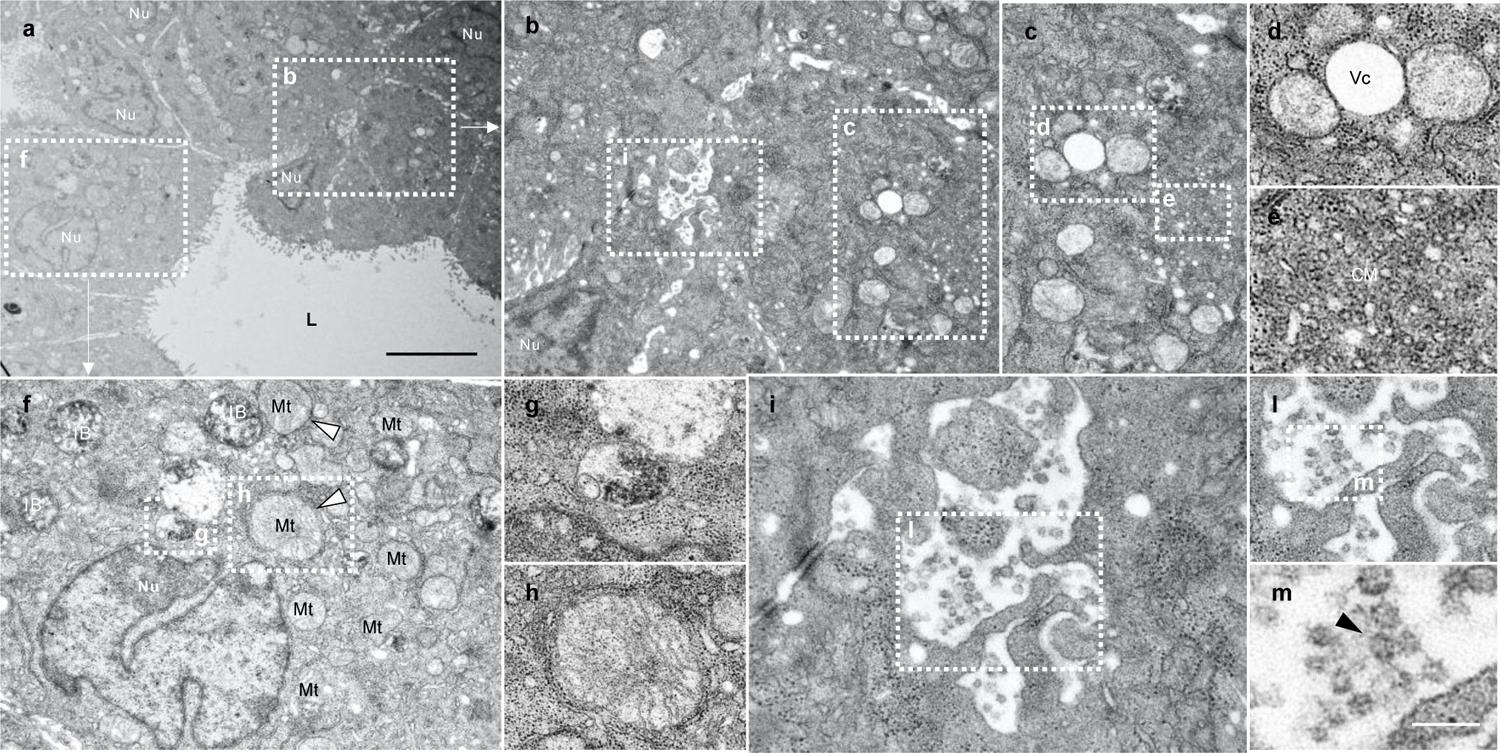
Transmission electron microscopy of SARS-CoV-2 infected CDCA-treated GCOs. (**a**) Electron micrograph of CDCA-treated GCOs infected by SARS-CoV-2 (**b-m**). Different magnifications of (**a**). Areas magnified in individual panels are marked with dotted lines and the corresponding panel letter is shown in white. Key morphological features consistent with viral infection and apoptosis are shown, such as formation of pathologic vacuoles (Vc) (**b-d**); convoluted membrane (CM) **(e),** inclusion bodies (IB) (**f-g**) and swollen mitochondria (white arrowhead) (**f, h**). Micrographs (**i-n**) showing virions with visible nucleocapsid proteins (black arrowhead) released in the extracellular space. Black scale bar 5 μm, white scale bar 200 nm. Nu, nuclei; L, lumen; Vc, vacuoles; Mt, mitochondria; CM, convoluted membrane; IB, inclusion bodies.

**Extended Data Figure 6.**
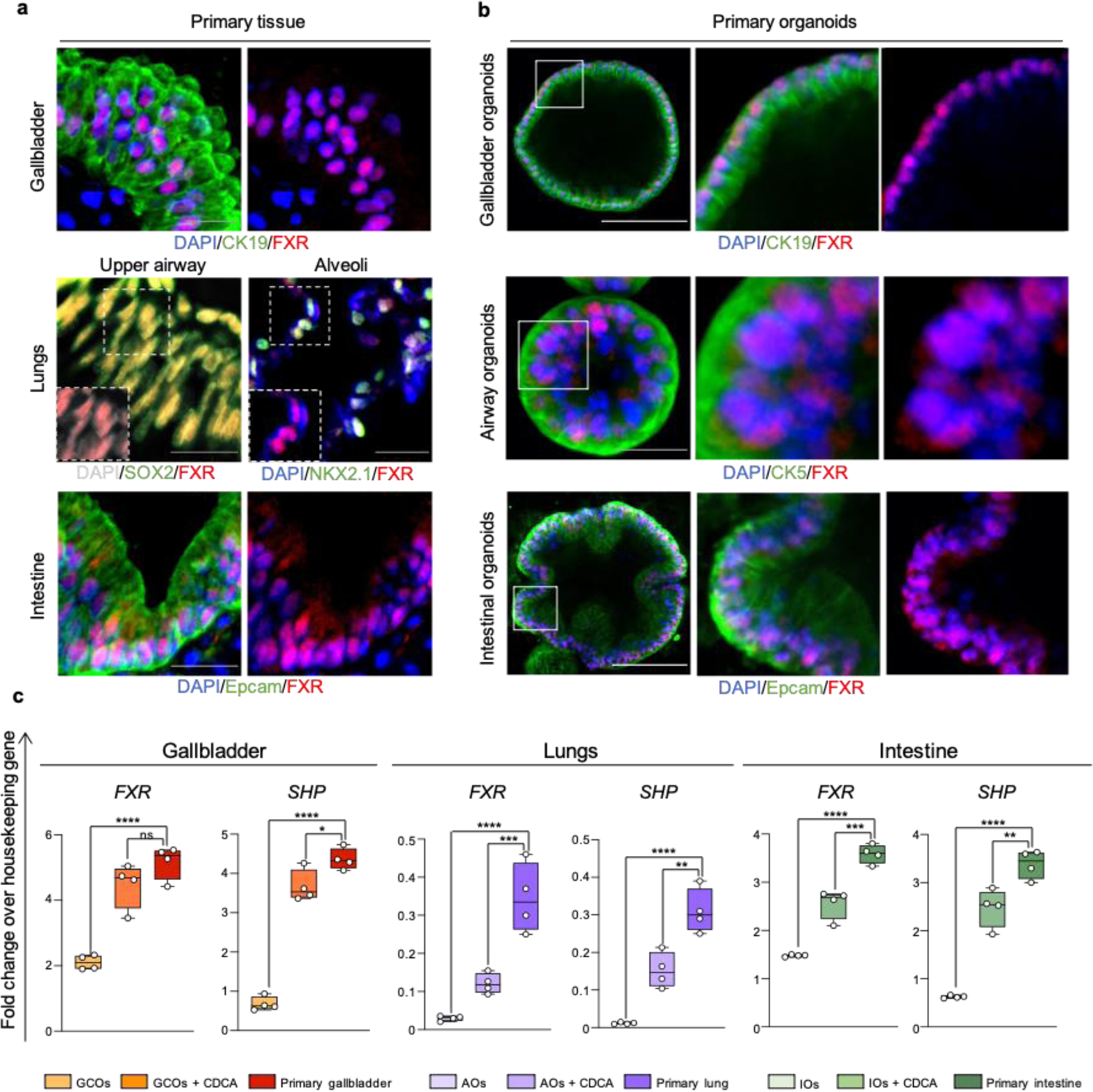
FXR is present and active in tissues affected by COVID-19 and their corresponding CDCA-treated organoids. (**a-b**) Immunofluorescence images confirming expression of FXR in primary human gallbladder, lungs and intestine (**a**) and in corresponding primary organoids treated with physiological levels of bile acids (CDCA) (**b**). White scale bars 100 μm; grey scale bar 25 μm. (**c**) QPCR analysis validating expression of FXR and its downstream effector SHP in primary tissue and corresponding organoids in presence or absence of physiological levels of bile acids (CDCA). CDCA treatment increases FXR and SHP expression in organoids to levels that are closer to primary tissue. Housekeeping gene, HMBS; n=4 independent samples. One-way ANOVA with Dunnett’s correction for multiple comparisons; ns, non-significant; **, *P*<0.005; ***, *P*<0.0005; ****, *P*<0.0001; center line, median; box, interquartile range (IQR); whiskers, range; error bars, standard deviation. GCOs, gallbladder cholangiocyte organoids; AOs, airway organoids; IOs, intestinal organoids.

**Extended Data Figure 7.**
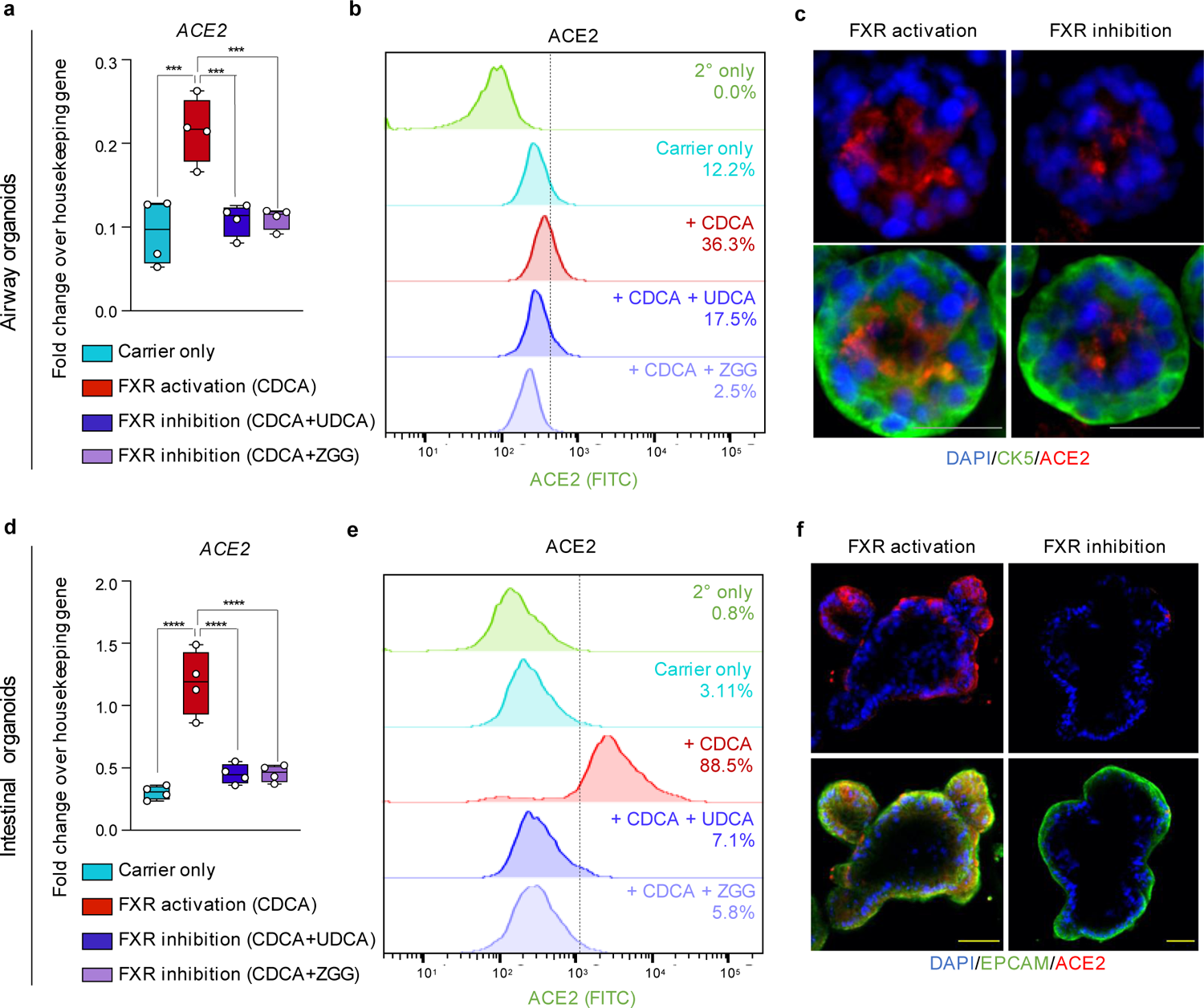
FXR regulates ACE2 in airway organoids and intestinal organoids. (**a-f**) QPCR (**a,d**), flow cytometry (**b,e**) and immunofluorescence (**c,f**) showing changes in ACE2 levels upon modulation of FXR activity in airway organoids (**a-c**) and intestinal organoids (**d-f**). Housekeeping gene, HMBS; n=4 independent samples. One-way ANOVA with Dunnett’s correction for multiple comparisons; ns, non-significant; ***, *P*<0.0005; ****, *P*<0.0001; center line, median; box, interquartile range (IQR); whiskers, range; error bars, standard deviation. Scale bars 25 μm.

**Extended Data Figure 8.**
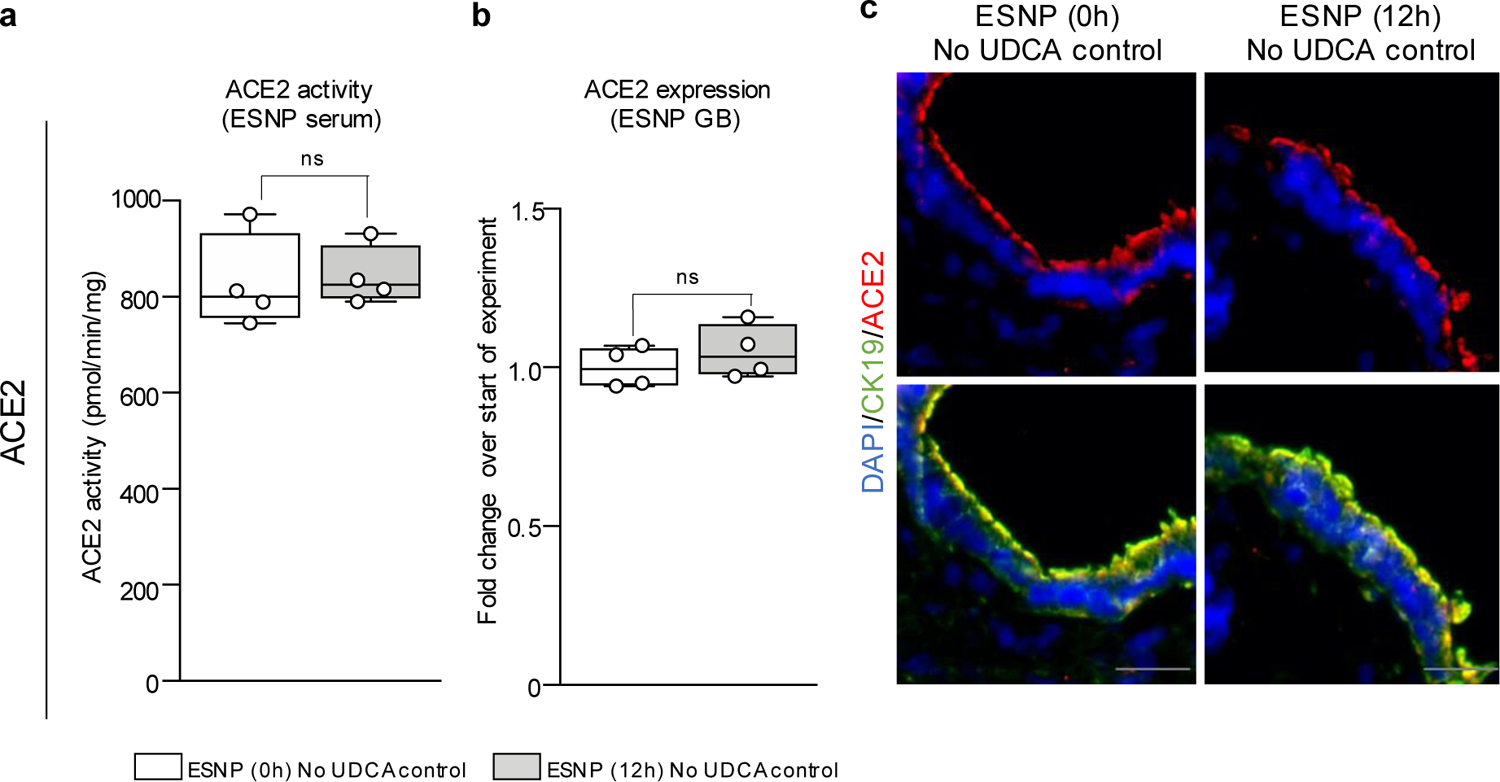
ACE2 levels are not altered by ESNP. (**a-c**) ACE2 enzymatic activity (**a**), QPCR (**b**) and immunofluorescence (**c**) showing that ESNP alone does not affect ACE2 levels. Housekeeping gene, HMBS; n=4 samples. Unpaired two-tailed t-test; ns, non-significant; center line, median; box, interquartile range (IQR); whiskers, range; error bars, standard deviation. Scale bars 25 μm.

