## Supplementary material for "FXR inhibition reduces ACE2 expression, SARS-CoV-2 infection and may improve COVID-19 outcome": Brevini et al., Supplementary Information

**This PDF file includes:**

**Supplementary Figure S1.** Gating strategy for flow cytometry analyses.

**Supplementary Table S1.** List of primary and secondary antibodies.

**Supplementary Table S2.** List of primers used for QPCR.

**Supplementary Table S3.** List of primers used for ChIP-QPCR.

**Supplementary Table S4.** Patient characteristics relative to 4f-h.

**Supplementary Table S5.** COVID-Hep and SECURE-Liver registries patient cohort characteristics.

**Supplementary Figure S1. Gating strategy for flow cytometry analyses**

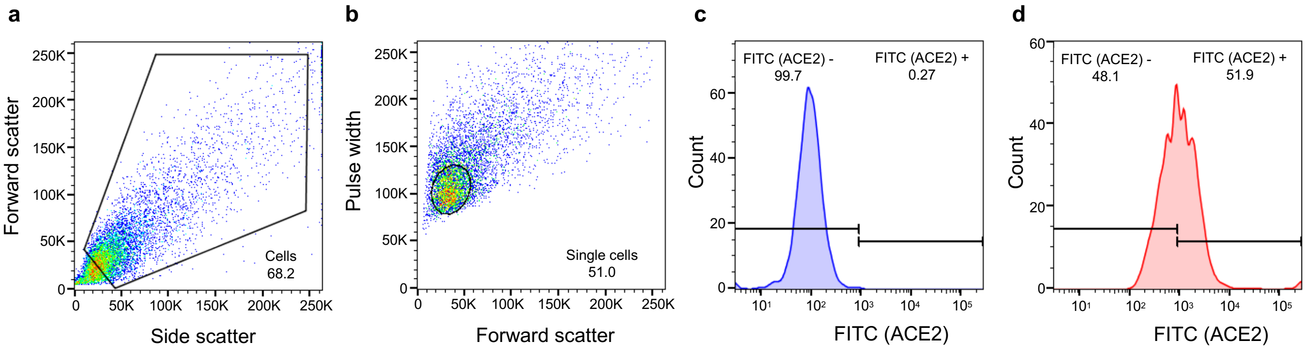


**Supplementary Figure S1. Gating strategy for flow cytometry analyses.** (**a-d**) Representative flow cytometry plots showing gating strategy for all flow cytometric analyses, gating on: (**a**) exclusion of debris; (**b**) exclusion of doublets; (**c**) secondary-only control to exclude negative population; (**d**) representative ACE2+ population.

**Supplementary Table S1.** List of primary and secondary antibodies

| Target protein | Company | Product code | Dilution |
|---|---|---|---|
| ACE2 | R&D | AF933 | 1:50 / 1:100 |
| ACE2 | abcam | ab15348 | 1:500 |
| ACE2 | abcam | ab108209 | 1:500 / 1:100 |
| EPCAM | R&D | MAB9601 | 1:50 / 1:100 |
| EPCAM | R&D | AF960 | 1:100 |
| Cytokeratin 19 | abcam | ab7754 | 1:100 |
| Cytokeratin 19 | abcam | ab52625 | 1:100 |
| SOX2 | abcam | ab15830 | 1:100 |
| SOX2 | R&D | AF2018 | 1:100 |
| NKX2.1 | abcam | ab72876 | 1:100 |
| Cytokeratin 5 | Thermo Fisher | MA5-17057 | 1:100 |
| SARS-CoV spike glycoprotein | abcam | ab273433 | 1:100 |
| SARS-CoV-2 nucleocapsid | Sino Biological | 40143-R019 | 1:100 |
| SOX17 | R&D | AF1924 | 1:100 |
| FXR | Novus biological | NBP2-16550 | 1:100 |
| FXR | Santa Cruz | sc-25309 X | 1:100 |
| Alexa Fluor Donkey Anti-Rabbit 568 | Invitrogen | A10042 | 1:1000 |
| Alexa Fluor Donkey Anti-Rabbit 488 | Invitrogen | A21206 | 1:1000 |
| Alexa Fluor Donkey Anti-Rabbit 568 | Invitrogen | A10042 | 1:1000 |
| Alexa Fluor Donkey Anti-Goat 488 | Invitrogen | A11055 | 1:1000 |
| Alexa Fluor Donkey Anti-Goat 568 | Invitrogen | A11057 | 1:1000 |
| Alexa Fluor Donkey Anti-Mouse 488 | Invitrogen | A21202 | 1:1000 |
| Alexa Fluor Donkey Anti-Mouse 647 | Invitrogen | A31571 | 1:1000 |

**Supplementary Table S2.** List of primers used for QPCR

| Target gene | Primer sequence (5' – 3') | |
|---|---|---|
| *HMBS* | Forward | GGAGCCATGTCTGGTAACGG |
| | Reverse | CCACGCGAATCACTCTCATCT |
| *GAPDH* | Forward | AGGACTCATGACCACAGTCCATGC |
| | Reverse | GATGACCTTGCCCACAGCCTT |
| *KRT19* | Forward | ACGACCATCCAGGACCTGCGG |
| | Reverse | TCCCACTTGGCCCCTCAGCGTA |
| *ACE2* | Forward | CTCCTAACCAGCCCCCTGTT |
| | Reverse | TGGAGGCATAAGGATTTTCTCCAC |
| *SARS-CoV-2 RdRp* | Forward | ATGGGTTGGGATTATCCTAAATGTGA |
| | Reverse | GCAGTTGTGGCATCTCCTGATGAG |
| *MS2* | Forward | TGGCACTACCCCTCTCCGTATTC |
| | Reverse | GTACGGGCGACCCCACGATGAC |
| *NRB02* | Forward | CCTGCCTGAAAGGGACCATCC |
| | Reverse | GCACCAGGGTTCCAGGACTTC |
| *NR1H4* | Forward | GCTTTGCTGAAAGGGTCTGC |
| | Reverse | CAGAATGCCCAGACGGAAGT |
| *IL-1β* | Forward | AAACAGATGAAGTGCTCCTTCCAGG |
| | Reverse | TGGAGAACACCACTTGTTGCTCCA |
| *IL-6* | Forward | AATTCGGTACATCCTCGACGG |
| | Reverse | GGTTGTTTTCTGCCAGTGCC |
| *IFNα* | Forward | GACTCCATCTTGGCTGTGA |
| | Reverse | TGATTTCTGCTCTGACAACCT |
| *IFNλ* | Forward | TCGCTTCTGCTGAAGGACTGCA |
| | Reverse | CCTCCAGAACCTTCAGCGTCAG |
| *KRT7* | Forward | GATTGCTGGCCTTCGGGGT |
| | Reverse | TCATCACAGAGATATTCACGGCTC |
| *CFTR* | Forward | AGTTGCAGATGAGGTTGGGC |
| | Reverse | AAAGAGCTTCACCCTGTCGG |
| *GGT1* | Forward | GTGAGAGCAGTTGGCTGTGC |
| | Reverse | GTTGAACTCTGCTGTGGGGC |
| *SOX17* | Forward | CGCACGGAATTTGAACAGTA |
| | Reverse | GGATCAGGGACCTGTCACAC |

*KRT19*, CK19; *NRB02*, SHP; *NR1H4*, FXR; *KRT7*, CK7.

**Supplementary Table S3.** List of primers used for ChIP-QPCR

| Target gene | Primer sequence (5' – 3') | |
|---|---|---|
| *OST$\alpha$* | Forward | AGTTCAGGGCTTTGGGTAATTAAAC |
| | Reverse | GGTGGAGGTCAGGGAAGGAAGA |
| *ACE2* | Forward | CGCTATCTTGAGGAAGAAGGGGAA |
| | Reverse | AGCAGGTACAAAGCATATGCAACC |
| *ACE2* negative region | Forward | AAGCGAGCTCAGTGTCCTCA |
| | Reverse | AGGTAGGCCCTTGAACCCTG |

**Supplementary Table S4.** Patient characteristics relative to 4f-h.

| | Age | Sex | Diagnosis | UDCA |
|---|---|---|---|---|
| Relative to 4f | 56 | F | Liver transplantation, cholangiopathy | Yes |
| | 33 | F | Gallstones/ microlithiasis | Yes |
| | 90 | F | Gallstones/ microlithiasis | Yes |
| | 74 | M | Liver transplantation - waiting elective procedure | No |
| | 74 | M | Liver transplantation - waiting elective procedure | No |
| | 40 | F | Liver transplantation - waiting elective procedure | No |
| Relative to 4h | 55 | F | Primary Biliary Cholangitis | No |
| | 71 | F | Primary Biliary Cholangitis | No |
| | 81 | F | Primary Biliary Cholangitis | No |
| | 56 | F | Primary Biliary Cholangitis | No |
| | 67 | F | Primary Biliary Cholangitis | No |
| | 64 | F | Primary Biliary Cholangitis | No |
| | 72 | F | Primary Biliary Cholangitis | No |
| | 52 | F | Primary Biliary Cholangitis | No |
| | 65 | F | Primary Biliary Cholangitis | No |
| | 43 | F | Primary Biliary Cholangitis | No |
| | 77 | F | Primary Biliary Cholangitis | No |
| | 71 | F | Primary Biliary Cholangitis | Yes |
| | 64 | F | Primary Biliary Cholangitis | Yes |
| | 71 | F | Primary Biliary Cholangitis | Yes |
| | 49 | F | Primary Biliary Cholangitis | Yes |
| | 50 | F | Primary Biliary Cholangitis | Yes |
| | 59 | F | Primary Biliary Cholangitis | Yes |

UDCA, ursodeoxycholic acid; F, female; M, male.

**Supplementary Table S5.** COVID-Hep and SECURE-Liver registries patient cohort characteristics

| | Total | No UDCA | UDCA | Propensity score matched (10:1) (No UDCA:UDCA) |
|---|---|---|---|---|
| **n, total** | 1096 | 1065 | 31 | 310 |
| **Sex (Female)** | 412 (37.6%) | 384 (36.1%) | 28 (90.3%) | 280 (90.3%) |
| **Age (years; median, IQR)** | 59 (48–68) | 59 (48–68) | 56 (44–66) | 57 (45–66) |
| ***Liver disease severity*** | | | | |
| **CLD, no cirrhosis** | 483 (44.1%) | 459 (43.1%) | 24 (77.4%) | 240 (77.4%) |
| **CTP A cirrhosis** | 272 (24.8%) | 270 (25.4%) | 2 (6.5%) | 2 (6.5%) |
| **CTP B cirrhosis** | 200 (18.3%) | 196 (18.4%) | 4 (12.9%) | 40 (12.9%) |
| **CTP C cirrhosis** | 141 (12.9%) | 140 (13.2%) | 1 (3.2%) | 10 (3.2%) |
| ***BMI*** | | | | |
| **<18.5** | 25 (2.3%) | 23 (2.2%) | 2 (6.5%) | 9 (2.9%) |
| **18.5-24.9** | 341 (31.1%) | 332 (31.2%) | 9 (29.0%) | 82 (26.5%) |
| **25-30** | 306 (27.9%) | 300 (28.2%) | 6 (19.4%) | 86 (27.7%) |
| **30-35** | 179 (16.3%) | 176 (16.5%) | 3 (9.7%) | 50 (16.1%) |
| **35-40** | 66 (6.0%) | 65 (6.1%) | 1 (3.2%) | 24 (7.7%) |
| **40+** | 51 (4.7%) | 50 (4.7%) | 1 (3.2%) | 27 (8.7%) |
| **Unknown** | 128 (11.7%) | 119 (11.2%) | 9 (29.0%) | 32 (10.3%) |
| ***COVID-19 outcome*** | | | | |
| **Hospitalised** | 898 (81.9%) | 887 (83.3%) | 11 (35.5%) | 236 (76.1%) |
| **ICU requirement** | 332 (30.3%) | 331 (31.1%) | 1 (3.2%) | 58 (18.7%) |
| **ICU admission** | 240 (21.9%) | 239 (22.4%) | 1 (3.2%) | 39 (12.6%) |
| **Invasive ventilation** | 168 (15.3%) | 167 (15.7%) | 1 (3.2%) | 31 (10.0%) |
| **Death** | 220 (20.1%) | 219 (20.6%) | 1 (3.2%) | 36 (11.6%) |

UDCA, ursodeoxycholic acid; IQR, interquartile range; CLD, chronic liver disease; CTP, *Child-Turcotte-Pugh* class; BMI, body mass index; ICU, intensive care unit.
